# Single cell transcriptomics identifies conserved regulators of neurosecretory lineages

**DOI:** 10.1101/2022.05.11.491463

**Authors:** Julia Steger, Alison G. Cole, Andreas Denner, Tatiana Lebedeva, Grigory Genikhovich, Alexander Ries, Robert Reischl, Elisabeth Taudes, Mark Lassnig, Ulrich Technau

## Abstract

Communication in bilaterian nervous systems is mediated by electrical and secreted signals, however, the evolutionary origin and relation of neurons to other secretory cell types has not been elucidated. Here we use developmental single cell RNA-sequencing in the cnidarian *Nematostella vectensis*, representing an early evolutionary lineage with a simple nervous system. Validated by transgenics, we demonstrate that neurons, stinging cells, and gland cells arise from a common multipotent progenitor population. We identify the conserved transcription factor gene *SoxC* as a key upstream regulator of all neurosecretory lineages and demonstrate that *SoxC* knockdown eliminates both neuronal and secretory cell types. While in vertebrates and many other bilaterians neurogenesis is largely restricted to early developmental stages, we show that in the sea anemone differentiation of neurosecretory cells is maintained throughout all life stages, and follows the same molecular trajectories from embryo to adulthood, ensuring lifelong homeostasis of neurosecretory cell lineages.

## INTRODUCTION

Metazoan cell type diversity is thought to have originated from multifunctional ancestral cell types which underwent segregation and subfunctionalization (Wagner, 2021). Driven by genetic individuation, descending sister cell types diverge and adopt new functions, which is pivotal for key innovations like a complex nervous system (Arendt et al., 2016; Murphy et al., 2019). While the evolutionary origin of neurons is still debated, it is known that the existence of chemical signaling predated the emergence of the neurons and therefore synaptic transmission (Burkhardt & Jékely, 2021). These observations suggest that the first form of cell communication was the secretion of neuropeptides (Jékely, 2021; Robitzki et al., 1989; Schuchert, 1993; Skorokhod et al., 1999). Recent advancements in genomics and single-cell transcriptomics provide opportunities to investigate the character of early neurosecretory cells and their relationships to other cell types, based on unbiased evaluation of gene usage across cells and organisms (Grundfest, 1959; Horridge, 1968; Mackie, 1990; Musser et al., 2019; Sachkova et al., 2021; Tarashansky et al., 2021).

In order to trace the evolution of neural cell types and nervous systems, it is most promising to investigate early branching metazoans. Ctenophores and Cnidaria are the first metazoan phyla in evolution with a nervous system. While the evolutionary relationship of ctenophores (and of their neurons) among animals is still contentious (Jékely & Budd, 2021; Kapli et al., 2021; Whelan et al., 2017), Cnidaria form the sister group to the Bilateria and are therefore highly informative regarding the evolutionary origin and diversification of bilaterian neural cell types. In the hydrozoan *Hydra*, the interstitial stem cell lineage gives rise to all differentiated cell types except epithelial cells, i.e. neurons, cnidocytes (also called nematocytes), gland cells and gametes (Bosch, 2009). In some species, e.g. *Hydractinia echinata*, interstitial stem cells are even considered totipotent (Frank et al., 2009). However, interstitial stem cells have so far only been identified in hydrozoans, which also possess several other derived features, and therefore we set out to study neurosecretory cell type relationships in the sea anemone *Nematostella vectensis*. Sea anemones belong to the Anthozoa among Cnidaria and are considered to have preserved more ancestral traits than hydrozoans (Bridge et al., 1992; Technau et al., 2005). The sequenced genome and recent chromosomal assembly of *Nematostella* have revealed a gene content similar to bilaterians, thus making it a phylogenetically highly relevant model system to investigate cell type evolution (Putnam et al., 2007; Zimmermann et al., 2020).

The nervous system of *Nematostella* includes sensory and ganglion neurons, which form diffuse neuronal networks in both cell layers (Nakanishi et al., 2012). The phylum-defining stinging cells (cnidocytes) are commonly regarded as the sister cell type of neurons, since both originate from a common neural progenitor population (Richards & Rentzsch, 2014). Furthermore, functionally diverse gland and muscle cell populations are present (Babonis et al., 2019; Cole et al., 2020; Sebé-Pedrós et al., 2018). Together, neurons, cnidocytes and gland cells form the neurosecretory lineages in cnidarians. These cell types are of particular interest, because of their vital functions for the organism, but also because they are terminally differentiated and must be replenished in a growing and dynamic organism. Indeed, in *Hydra*, epithelial cells divide approximately every 4 days under regular feeding conditions, meaning that the nervous system would be diluted by half within 4 days if not supplemented by newly differentiating neurons from stem cells. Moreover, cnidocytes are discharged and destroyed during feeding, and therefore must be constantly replaced. While recent work has revealed several conserved transcription factors and signaling pathways involved in neurogenesis during embryonic and larval stages, it is currently unclear whether these processes continue to occur in adult polyps during conditions of homeostasis, or whether different mechanisms specific to homeostasis are implemented.

In order to determine the developmental origins and relationships of neurosecretory cell types, we generated a single-cell RNA-seq dataset, which integrates 11 time points from embryogenesis to the adult polyp. Our data suggest that neurons share a closer relationship with gland cells than with cnidocytes. We identified several regulatory factors marking cell fate decisions underlying the replacement of cell types throughout homeostasis and validated their predicted roles at key developmental decision points in transgenic lines. Reconstruction of the developmental pathways within the neurosecretory partition revealed a common progenitor population, which gives rise to all neurosecretory cell types during embryogenesis and is maintained throughout homeostasis. These multipotent progenitors are specified by a hierarchy of *Sox* transcription factors and we identify *SoxC* as an upstream regulator of *SoxB2a*, a key neuronal transcription factor in vertebrate development (Sandberg et al., 2005). Interestingly, Sox genes are drivers of cell fate decisions in the vertebrate neural crest, and members of the *SoxC* family regulate the acquisition of neural progenitor identity and subsequent differentiation (Kavyanifar et al., 2018), suggesting an ancestral role of this transcription factor family in the diversification of neurosecretory cell lineages.

## RESULTS

### A developmental single-cell atlas of *Nematostella vectensis*

To trace the appearance, developmental origins, and lineage relationships between all neurosecretory cell types, we generated a time-course of single-cell transcriptomic libraries (**Fig. 1A**). We included developmental stages from the blastula-gastrula transition (18 hpf), mid-late gastrula (24 and 25 hpf), a series of planula larval stages (2, 3, 4 dpf), the planula-polyp transition (5 dpf), and primary polyp (8 and 16 dpf) stages. This developmental series was complemented by a set of 6 tissue-specific libraries that provide spatial information: Tentacles, body wall, pharynx, pharynx with body wall, and two mesenteries, which are foldings of mixed embryological origins within the gastric cavity (Steinmetz et al., 2017). A detailed summary of all sampled time-points and other specifications of the resulting 16 individual libraries are provided in **Fig. S2A**, all single-stage and tissue-specific libraries are supplied in **Fig. S1**. The entire dataset contains 55 042 cells after filtering (see methods), wherein each cell is represented by a median of 739 genes/cell and a median of 1782 nUMIS/cell. 85% of the detected genes are represented by more than 10 reads in our dataset, and the average gene model detection throughout all single libraries is 75% **(Fig. S2A)**, resulting in a detection of 97% of all gene models in the merged dataset. This surpasses the previously published single cell atlas (Sebé-Pedrós et al., 2018) both in terms of cell numbers as well as stages, allowing us to carry out detailed analyses regarding the molecular regulation underlying cell fate specification throughout the life cycle of *Nematostella*.

**Figure 1:**
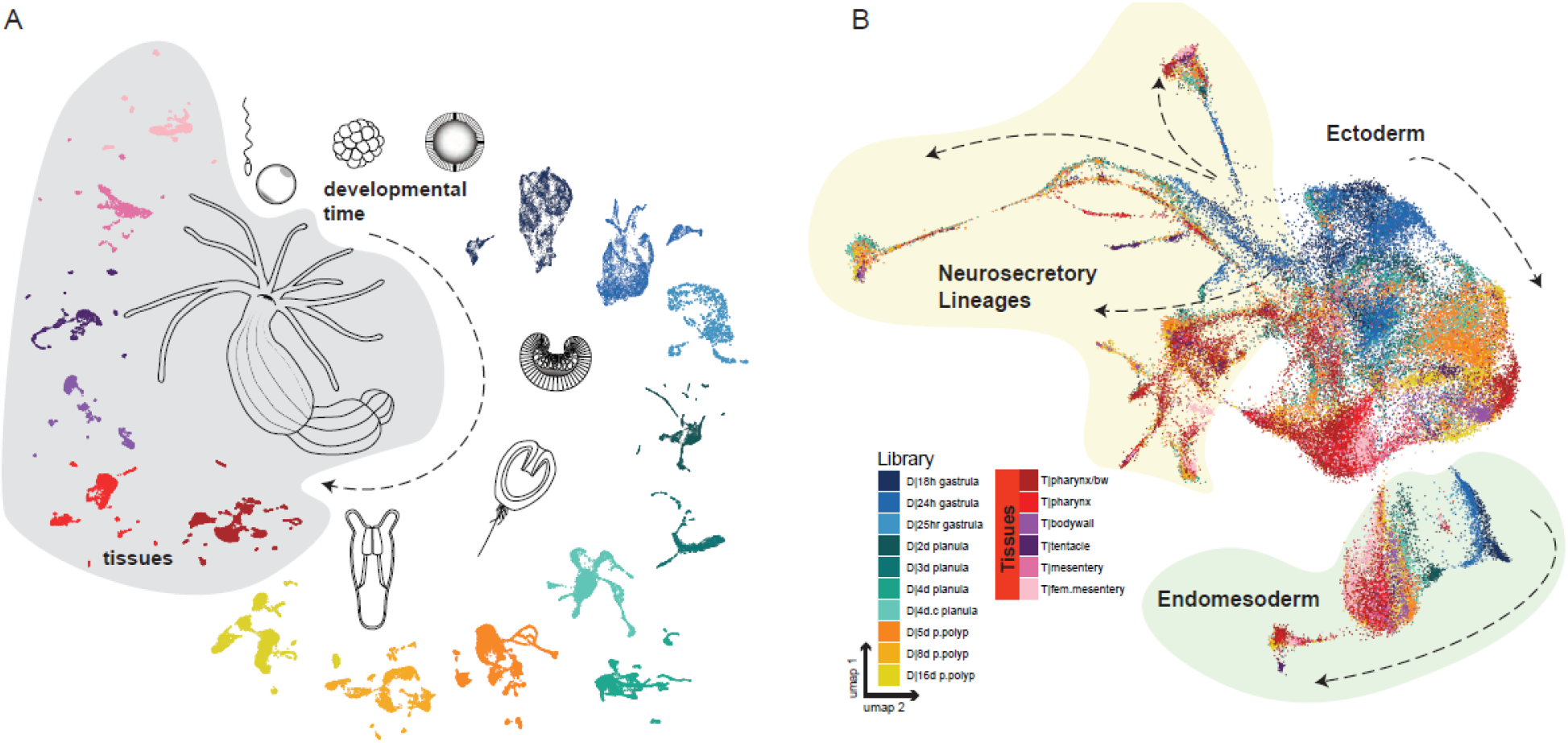
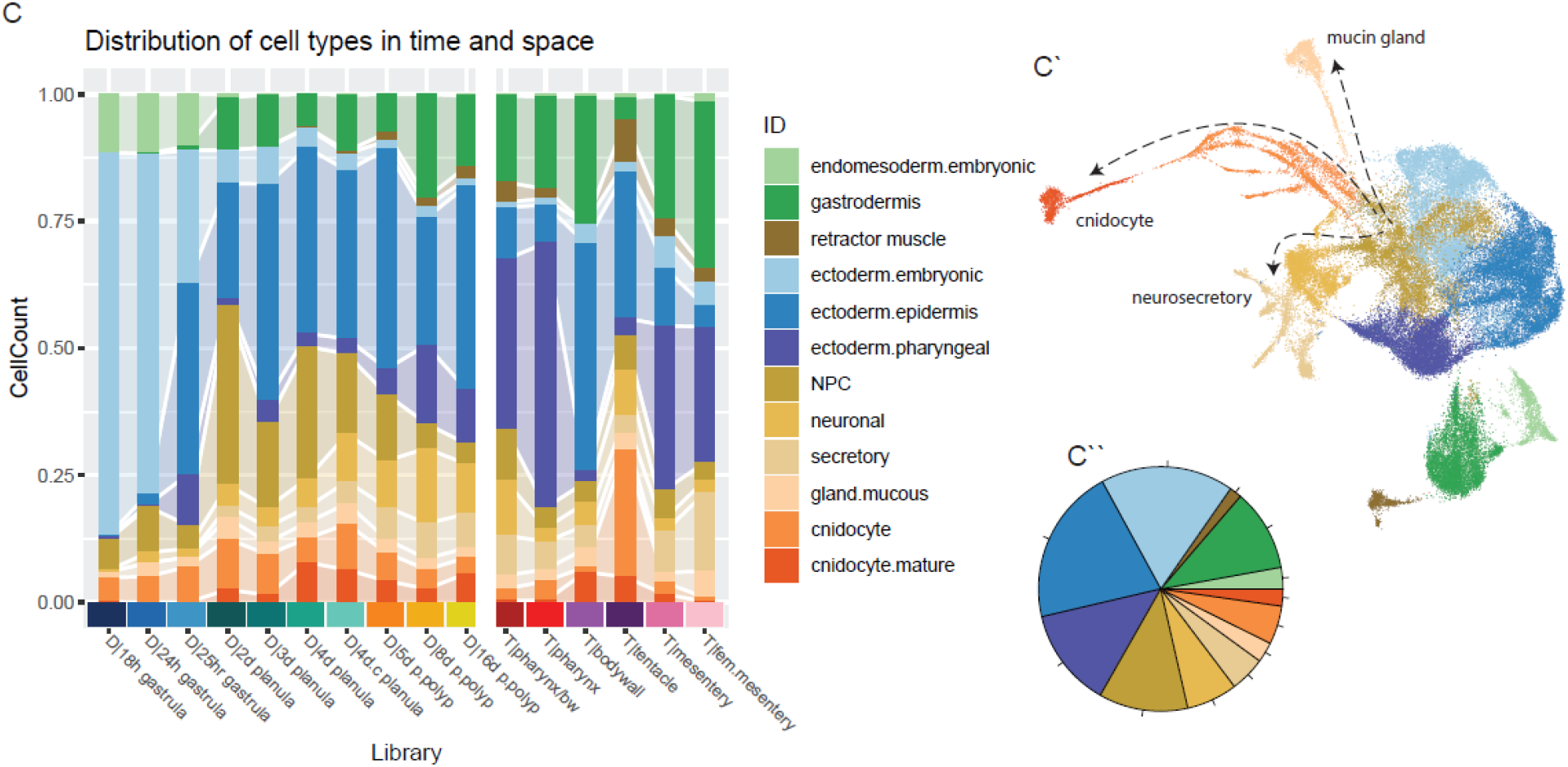
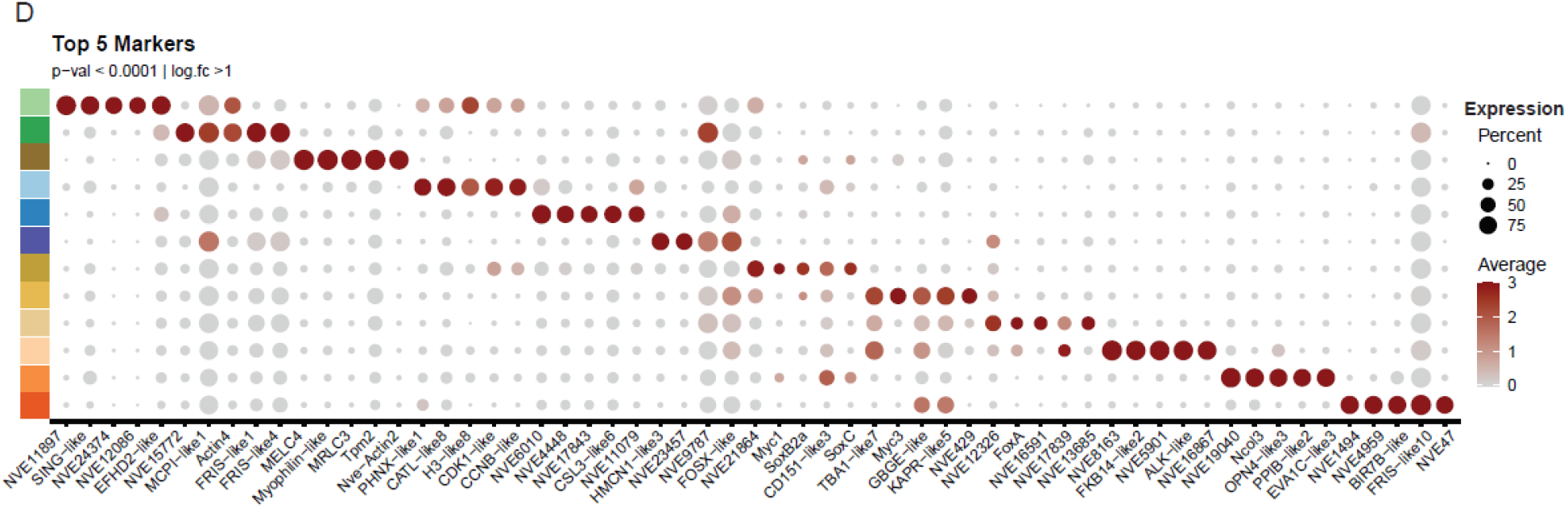
Single cell sequencing provides a transcriptomic map of development for the sea anemone Nematostella vectensis. A) 11 libraries from sequential developmental stages ranging from 18hpf to 16dpf (developmental series) and 6 tissue-specific libraries from tissues harvested from a sub-adult animal (tissues) were generated with the 10x Genomics platform. B) UMAP projections of the combined dataset illustrate partitioning of all cells into three distinct partitions: endomesoderm (green), neurosecretory cells (yellow), and ectodermal (white). Cells are colour-coded according to the library of origin: blue - yellow (developmental series) and red-purple (tissues), and clearly sort along the axis of time (dashed arrows). C) Clustering of the cells resolves into 12 populations, the proportion of which within each library is shown as a histogram. Three principle neurosecretory lineages emerge from the ectoderm (arrows). C’ UMAP projection color-coded by clusters. C’’ Distribution of populations across the entire dataset. D) Expression profile of top 5 marker genes for each cluster are shown as a dotplot; the size of the dot represents the percentage of cells in the population for which the gene is detected, and the color represents the relative expression values.

First, we integrated expression data from all libraries into a single dataset, clustered the dataset into broadly defined cell populations, and projected the data onto 2 dimensions using UMAP ((Becht et al., 2018), see **SI_1** for 3d projection with rotation). The entire dataset sorts into three broad partitions: endomesoderm (green), ectoderm (white), neurosecretory (yellow), each with a clearly evident temporal axis running from early (dark blue) to late (yellow) time points, terminating at the sub-adult tissue libraries (red-purple) (dashed arrows, **Fig. 1B**). Cell populations were annotated manually by combining analysis of the underlying molecular profile and prior knowledge (**Fig. S2B**). In total we defined 12 broad cell populations split across the three partitions (**Fig. 1C**), which are defined by distinct gene sets (**Fig. 1D, Table SI_2**). Two of these major groups corresponded to developmental stage-specific cell states of the two primary germ layers (“endomesoderm.embryonic”, “ectoderm.embryonic”. We recovered large populations corresponding to the mature gastrodermis and associated retractor muscle, outer ectodermal epithelium, and inner ectoderm (“ectoderm.pharyngeal” including septal filaments and pharyngeal ectoderm). Half of the cells in the whole dataset are assigned to the ectodermal populations (blue color shades), around 20% derive from the endomesoderm (green colors), and the remaining fraction constitutes the complement of neurosecretory cell types (shades of yellow/orange; **Fig. 1C’’**). We recover mucus-producing gland cells (“gland.mucous”), two cnidocyte populations (“cnidocyte”, “cnidocyte.mature”), and a large heterogeneous neurosecretory population (“neuronal”, “secretory”) (**Fig. 1C’)**. All of these populations are characterized by the presence of intermediate differentiation states, most prominently in the trajectory underlying cnidogenesis (orange shades **Fig. 1C’**). Strikingly, recovered developmental pathways predicted that all neurosecretory cells originate from a common progenitor population (ochre, **Fig. 1C’**). To explore this further, we focused on an in-depth investigation of the trajectories emerging from these putative neurosecretory progenitors and the relationships between their derivatives.

### Molecular regulation of cnidocyte specification, differentiation and maturation

Cnidocytes are a prominent apomorphic feature of cnidarians. They contain the cnidocyst, a dischargeable organelle, generated by accumulation and assembly of specific structural proteins in a post-Golgi vesicle (Özbek et al., 2009). Thanks to its toxin complement, this cell type is indispensable for defense and predation throughout the diverse phases of the life cycle. The recovered branching trajectory within the cnidocyte lineage likely reflects their high turnover rate and the corresponding increase in capture of cell states indicative of the specification processes (orange shades, dashed arrow “cnidocytes” **Fig. 1C’**). This suggests that the resolution of our dataset extends beyond the identification of cell types and contains a diversity of intermediate cell states, allowing us to reconstruct developmental trajectories. To test this idea, we extracted all cnidocyte-specific transcriptomes for a comprehensive characterization of the diversity of differentiation states. We also investigated the underlying regulatory signatures, which can shed light on the molecular requirements for the evolution of a novel cell type.

Previous analyses of the cnidom of *Nematostella* have suggested three morphologically distinct types of cnidocytes: spirocytes and two nematocyte types, the most abundant basitrichous haplonemas, which exist in two different sizes, and microbasic mastigophores (**Fig. 2A;** Zenkert et al., 2011, Babonis & Martindale, 2017; Karabulut et al., 2021). We were able to identify these three types in our dataset; the two basitrichous haplonemas of distinct sizes (also called isorhizas) correspond to the two most pronounced nematocyte trajectories (‘nem.1’ (green) and ‘nem.2’ (purple)). They share the largest overlapping transcriptomic profile, including a common toxin profile (**Fig. 2D**), but differ in their expression of several structural genes (in particular hemicentins and GAPR paralogs, **Fig. 2D, SI_2**). Additionally, the two branches have distinct distributions across our tissue-specific libraries, which enables an assignment of the two size variations of this nematocyte type, since large isorhizas are enriched in tentacles (Babonis & Martindale, 2017). Spirocytes (**Fig. 2B**; pink trajectory), which have a characteristic spiral thread in the capsule, appear in the 4d planula, are restricted to the tentacle tissue libraries, as is expected of this cell type, and express *FoxL2* (**Fig. 2E**, (Sebé-Pedrós et al., 2018)). The pharyngeal ectoderm harbors two additional cnidocyte populations, which may correspond to microbasic p-mastigophores (“cni.nep8”, “cni.unchar.ph”; **Fig. 2B**, (Matus et al., 2007)). Previous reports (Columbus-Shenkar et al., 2018; Moran et al., 2013) described cells exhibiting a gland-like cellular morphology within the pharyngeal ectoderm expressing *NEP8*. We identify these special cnidocytes as the subpopulation “cni.nep8” (yellow branch) expression *NEP8* together with *NEP6* and *NEP14*. A second, novel, pharynx-specific cnidocyte type (“cni.unchar.ph”, slate grey) expresses a specific set of uncharacterized genes suggesting this may be a *Nematostella*-specific cell-type related to the nem.2 haplonema lineage due to overlapping *aristaless* expression **(Fig. 2D,F)**. In addition, we identified cnidocytes with unique profiles that are stage specific. Cnidocytes within the gastrula (dark blue) have a distinct molecular profile and express a unique toxin, *TX60B-like3* (**Fig. 2C,D**). We further recovered three small populations, which are found exclusively in the planula larva (‘planula.nem’: dark orange; ‘cni.planula.spirocyte’: light blue; ‘planula.mature’: light orange). Of these, ‘planula.nem’ has a distinct toxin profile (*TX60B-like5* and *6, COMA-like1*, **Fig. 2C**), highlighting the dynamic and differential use of toxins throughout the life cycle (**Fig. 2B**).

**Figure 2.**
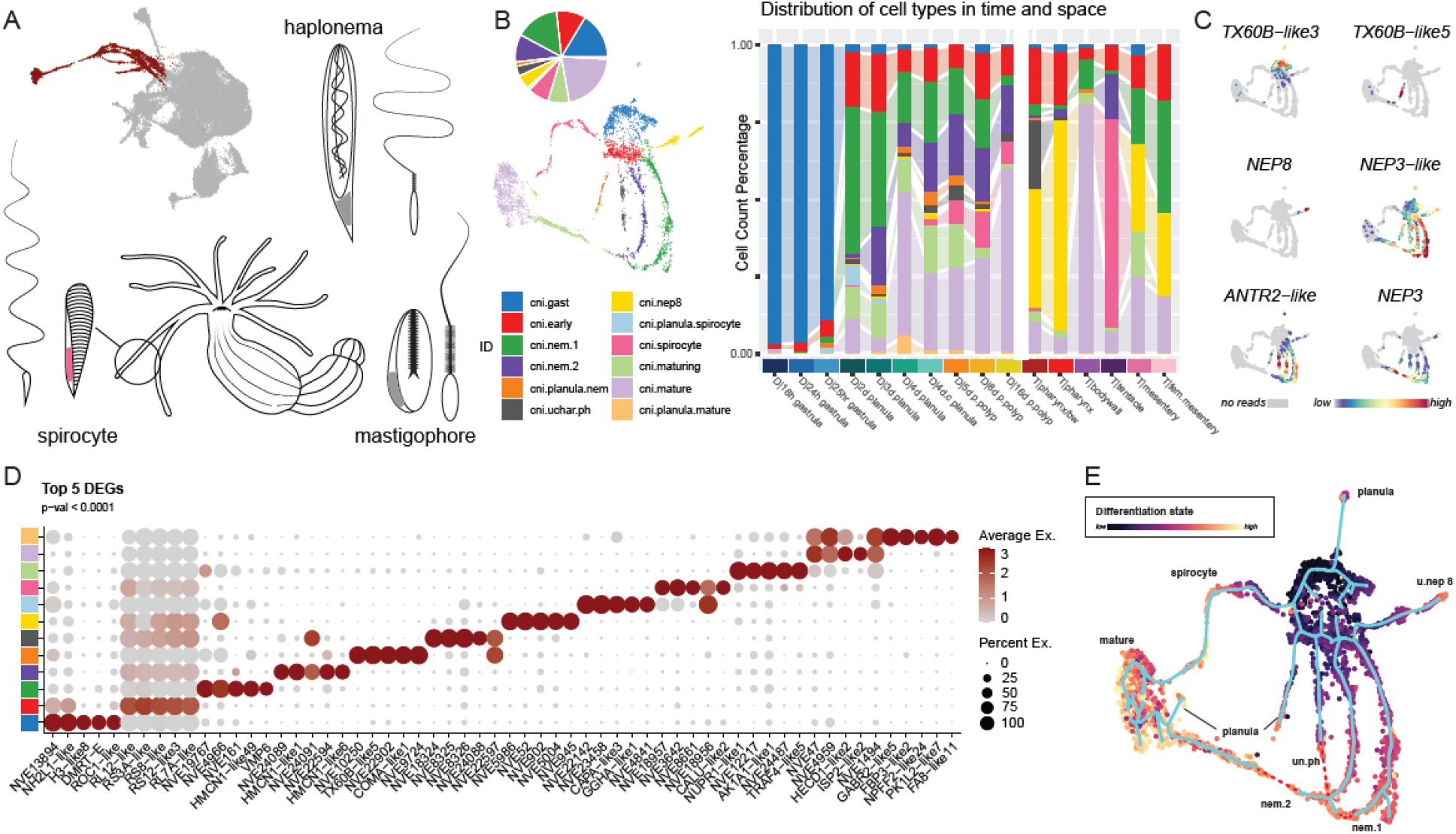
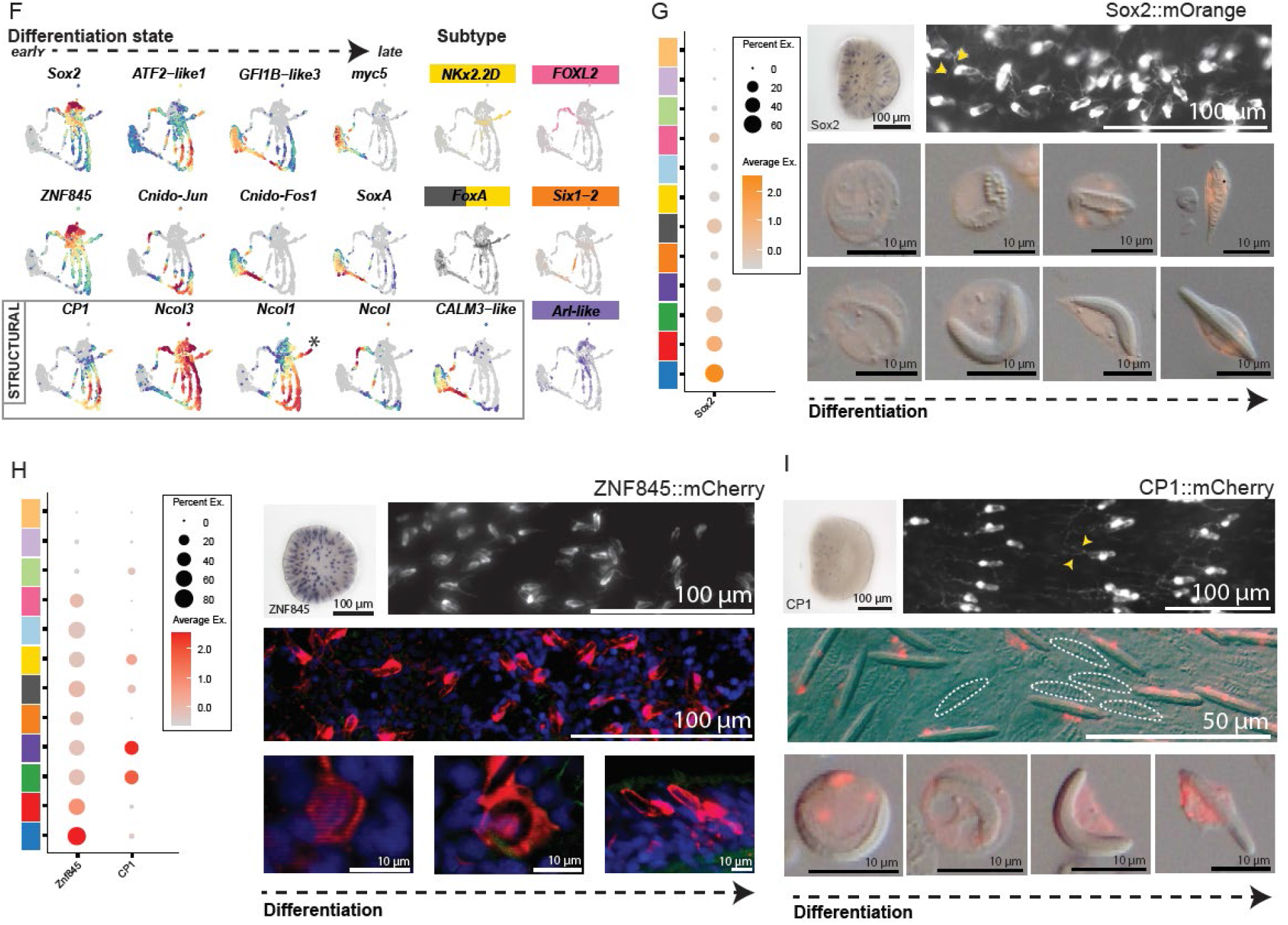
Analysis of the cnidocyte partition reveals convergence to a common transcriptomic phenotype across all cnidocyte cell types. A) Schematic representation of the three most prevalent cnidocyte subtypes previously described in Nematostella. (based on (Zenkert et al., 2011)). Cnidocyte partition of the dataset is highlighted in dark red (inset). B) Distribution of the identified unique transcriptomic profiles as proportion (pie), UMAP reconstruction of 4035 cnidocytes, and library contribution (barplot). C) Unique toxin profile of identified specification branches. D) Gene expression dotplot illustrating that each specification trajectory is supported by a set of differentially expressed genes. E) cytoTRACE inference of differentiation state shows maturation trajectories of each cnidocyte subtype from a centrally located common progenitor population. Reconstructed trajectories (Monocle 3) shown in blue. F) Gene expression profiles of regulatory molecules involved in the specification of the cnidocyte lineage (top 2 rows) and dynamic expression patterns of structural genes (bottom row). G,H,I) ZNF845::mCherry, Sox2::mOrange and CP1::mCherry transgenic lines confirm the specificity of gene expression profiles and visualize the morphology of cnidogenesis. Left panel: Dotplot of gene expression profile across the populations colored as in (B). In situ expression profiles show single cell gene expression within the ectoderm. Transgenic animals exhibit fluorophore expression with all cnidocyte populations under the promoter of ZNF845 and Sox2 (G, H), but absent from spirocytes when driven by the CP1 promoter (I: white dashed lines). Sequential developmental stages are identifiable in both lines from dissociated cell spreads.Axon-like processes of nematocytes are indicated by yellow arrowheads (G, I)

Cnidocytes are destroyed when they are used in predation or defense, and so are continuously replaced. We therefore presumed that we should be able to sample most cell states during the differentiation of various subtypes given this high turn-over rate. We predicted the differentiation state of all cells in the subset with CytoTRACE, which uses the number of expressed genes as a predictor of developmental potential (Gulati et al., 2020), and predicted cell trajectories using the learn_graph function in Monocle3 (Cao et al., 2019). Early cnidocyte specification factors identified include lineage-specific *Sox2*, the zinc-finger containing gene *ZNF845*, myc-family bHLH genes *Myc4* and *MAX-like*, and PRDM-family members *PRDM6-like* and *PRDM13* **(Fig. 2F; SI_2)**.

To confirm the expression of *Sox2* and *ZNF845* in early primed cnidocytes, we generated *Sox2::mOrange2* and ZNF845::mCherry transgenic lines and observed that the fluorophore reporter is expressed in differentiating haplonemas, already numerous in the gastrula **(Fig. 2G, E)**. In line with recent observations, these cells display neurite-like structures (Karabulut et al., 2021), a feature that has not been described for hydrozoan cnidocytes (**Fig. 2G;** yellow arrowheads). Furthermore, the stability of reporter protein allowed us to trace the morphological changes throughout differentiation, with a confident identification of *Sox2*-driven mOrange2+ cnidocytes within dissociated cell suspensions. Our Sox2 reporter line revealed that the capsule has a curved morphology, which straightens as differentiation proceeds, accompanied by an increased capsule to cytoplasm ratio. The maturation of cnidocytes is also visualized by mosaic transgenic ZNF845+ cells, which show the progression from small and curved capsule to fully differentiated nematocyte, arranged within the tentacle ectoderm **(Fig. 2E)**. Consistent with an early role in cnidocyte specification **(Fig. 2G)**, we detected *Sox2*-driven mOrange2 expression in both nematocytes and spirocytes **(Fig. 2G)**.

Our molecular characterization indicates that the transition to the mature cnidocyte phenotype in all lineages is characterized by sequential activation of specific members of the cFos/Jun and ATF/CREB pathways, and activation of the zinc-finger protein GFI1B-like3 (**Fig. 2F**). The temporal progression through differentiation via sequential expression of JUN/FOS paralogs has been partially reported previously (Sunagar et al., 2018). Knockdown of *Cnido-Jun* disrupts normal nematogenesis, which is consistent with expression in the early tract of our differentiation trajectory. *Cnido-Fos1* expression follows that of *Cnido-Jun*, which suggests a role related to the onset of maturation (Sunagar et al., 2018). This shift in transcription factor profile is accompanied by the expression of a cnidocyte-specific peptidase, *CP1*. To validate this finding, we also generated a transgenic reporter line using the P1 promotor (**Fig. 2I**). At the gastrula stage, *CP1* gene is already expressed in fewer cells than *Sox2*, indicating its lineage restriction **(Fig. 2I)**. Notably, spirocytes are negative for mOrange2 reporter expression. This suggests that this gene is either activated after the decision between spiro-or nematocyte trajectories or is rapidly repressed during spirocyte differentiation. Together with the reconstructed pseudotime trajectories, this suggests nematocytes and spirocytes share a common *Sox2+* progenitor and that the fate decision between those two lineages may either occur later or in combination with another factor.

Strikingly, all cnidocyte trajectories emerge from a single pool of progenitor cells (**Fig. 2B**; red population) and diverge into the distinct capsule types. Subsequently, the expression of subtype-specific structural genes ceases and a molecular switch causes all branches to converge upon a common transcriptomic profile which terminates in a single mature cell cluster (**Fig. 2E)**. This is a feature that to our knowledge has not been reported for other cell type differentiation trajectories. In our single-cell atlas, the terminal mature state shared by all different cnidocyte subtypes has a distinct regulatory signature, including two *myc* genes and *SoxA* (**Fig. 2F)**, and is also enriched in calcium-binding genes and ion channels, **(SI_2**, *Calbindin-32* (*CAB32-like*), calmodulins (*CALM3-like, CALM-like7*) and the calcium channel polycystine *PKD2-like4* (McLaughlin, 2017). Since cnidocyte discharge is a calcium-dependent exocytotic process, which can be triggered by chemosensory and mechanosensory stimuli (Balasubramanian et al., 2012; Özbek et al., 2009), we conclude that this final transcriptomic state is directly linked to the ability to discharge. The upregulation of calcium-binding protein genes is accompanied by a decrease in minicollagen gene expression at the beginning of the maturation phase **(Fig. 2F)**, which suggests that once all structural components of the different capsule types are formed, the transcriptome quickly adapts to its next and final function in discharge.

Cnidocytes are commonly considered as a sister cell type of neurons, a developmental relationship which is reflected in our full dataset (**Fig. 1C’**, Richards & Rentzsch, 2014). The highly resolved cnidocyte differentiation trajectory demonstrates that specification pathways can be reconstructed from our dataset. Therefore, we then extended our analysis to the entire neurosecretory partition, with the aim to understand the developmental origins and relationships of all lineages. We extracted neural progenitor cells (NPCs) together with all descending daughter cells and further separated this cell selection into three subsets: An embryonic subset, in order to study the onset of lineage specification during gastrula stages **(Fig. 3A**), which we expect to recapitulate what is known from previous studies on the earliest neurogenesis; from early planula to primary polyp stages **(Fig. 3C**) to determine whether these patterns are retained during development; and finally a sub-adult subset that consists of all tissue-derived libraries **(Fig. 3E)** to address whether similar cell trajectories are retained during homeostasis. For each subset, we reconstructed putative cell lineage trajectories using the learn_graph function in Monocle3 (Cao et al., 2019), and assessed the differentiation potential of all cells with cytoTRACE (Gulati et al., 2020) (**Fig. 3A, C, E**).

**Figure 3:**
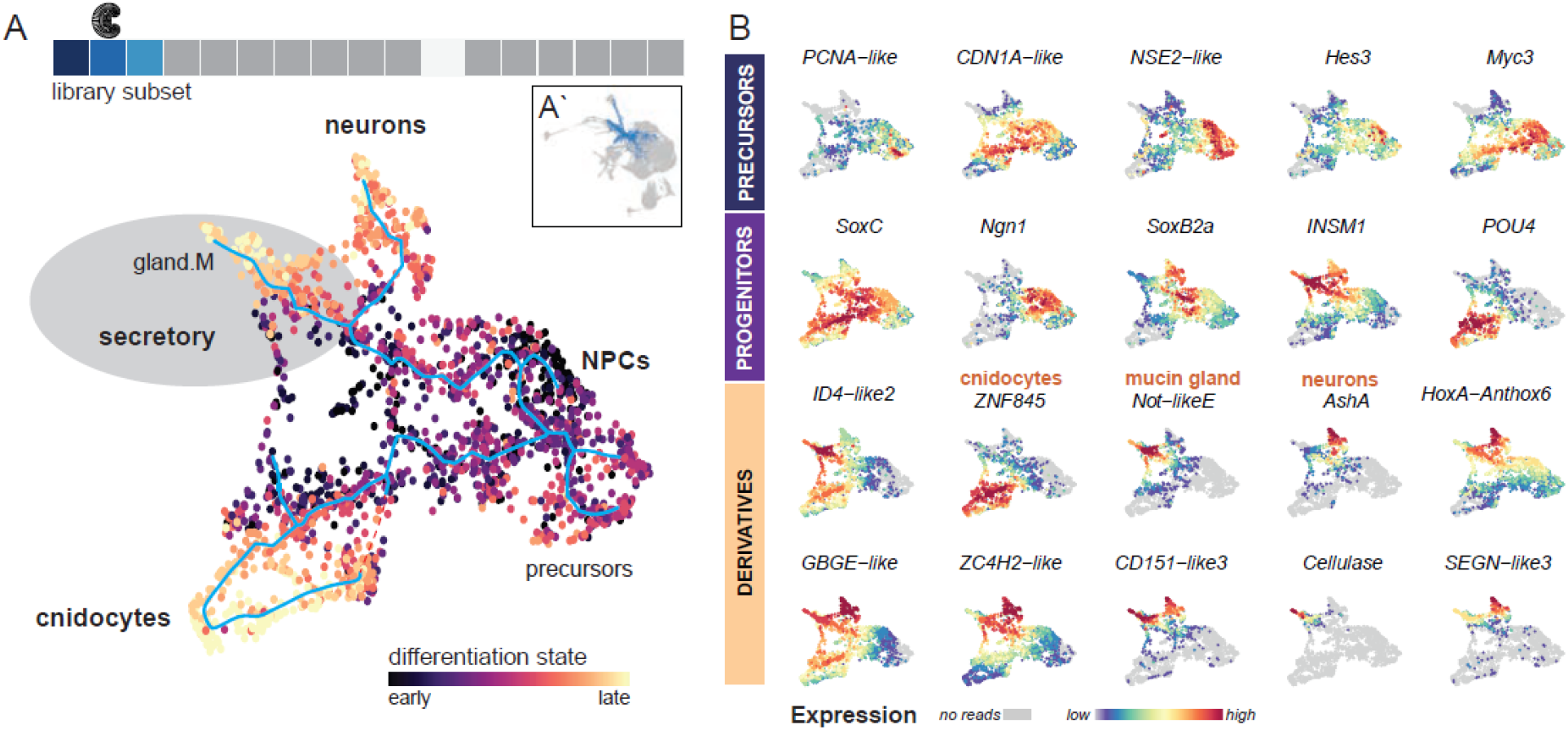
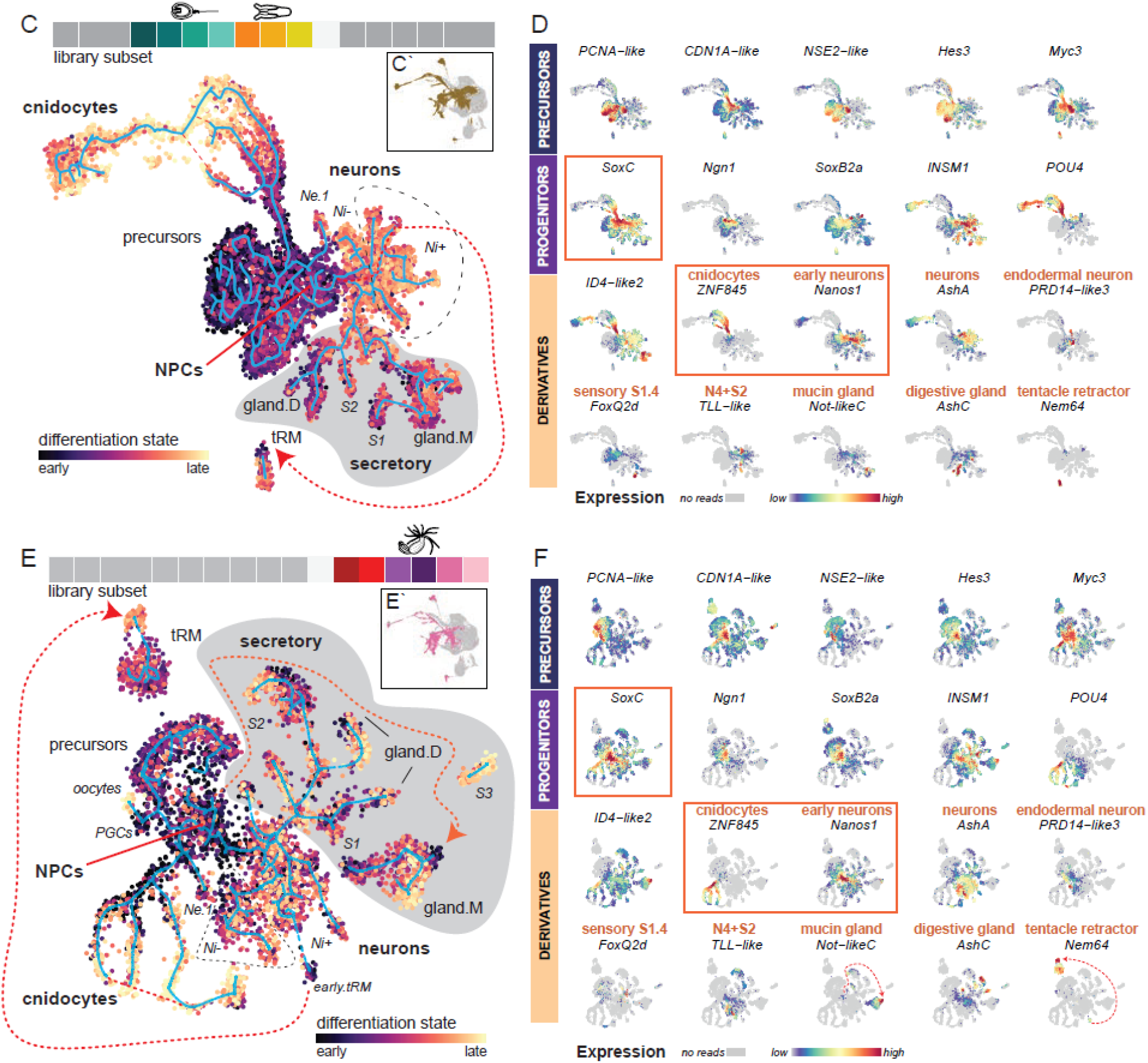
Analysis of entire neurosecretory lineages with progenitor cells reveals common molecular architecture of NPCs and robust differentiation trajectories during early development and homeostasis. A, C, E) Reconstructed trajectories (Monocle 3) shown in blue, cells are colour-coded according to predicted differentiation state with CytoTRACE. Dashed red lines indicate predicted relationships not recovered informatically. Bar at the top: Single library color code illustrates which developmental stages are included in the respective subset highlighted below. B, D, E) Gene expression patterns of representative gene models indicative of precursors (dark blue), progenitors (purple) and their derivatives (dark yellow). Orange boxes indicate expression profiles of genes selected for in vivo validation with transgenic lines in the scope of this paper.

With this approach we were able to reconstruct the previously reported cell type complement and lineage relationships, and extended this knowledge through identification of novel factors associated with early cell specification within the gastrula. Neurons (marked by *AshA;* (Layden et al., 2012)) and cnidocytes (marked by *ZNF845*, **Fig. 3B**) are derived from a *SoxB2a* progenitor cell population also known as *SoxB(2)* **(NPC; Fig. 3B** (Richards & Rentzsch, 2014, 2015). A third derivative of the NPCs arises as a bifurcation of the trajectory leading to the neuronal branch **(Fig. 3A)**. We identified these cells as mucin-producing gland cells (expressing *Mucin* (Steinmetz et al., 2017) and *Cellulase* (Sinigaglia C, 2015)) and found a distinct transcription factor signature consisting of two *notochord homeobox* paralogs *(Not-likeC* and *Not-likeE*, **Fig. 3B**). Derivatives of the NPCs are divided between *POU4* expressing cnidocytes and *Insulinoma1* (*Insm1*) expressing neurons and gland cells, which also express *HoxF/Anthox1*. Neurons and mucin gland cells share the expression of structural genes, such as the tetraspanin *CD151-like3* and the calcium-binding protein *SEGN-like3* **(Fig. S3T, U)**. Additionally, expression of the zinc finger transcription factor *ZC4H2-like* and the G-protein *GBGE-like* is detected in all three branches **(Fig. 3B)**.

Regulatory genes upstream of NPCs include the HMG-family member *Sox3*, zinc-finger protein *OZF-like*, E3 SUMO-protein ligase *NSE2-like*, and three paralogs of the bHLH family of transcriptional inhibitors of the HES family (“precursors”, **Fig. 3B**). These transcription factors are expressed in a population of cells with a low differentiation score, which is calculated based on the number of expressed genes per cell and thus predicts a high developmental potential of this cell population (**Fig. 3A**, Gulati et al., 2020). Furthermore, these undifferentiated cells exhibit pronounced cycling characteristics, indicated by two distinct molecular states reflective of a high rate of DNA synthesis and mitotic spindle organization, respectively. Based on these characteristics, these cells may represent the cycling pool of stem cells from which the NPC population emerges. Exit from mitotic cycling is accompanied by the expression of a cyclin-dependent kinase inhibitor (*CDN1A-like*/p21 **Fig. 3B**, (Xiong et al., 1993)). Downstream, we identify two *myc* orthologs, *Myc1* and *Myc3*, and the HMG-sox family regulator *SoxC*, whose expression is prominent along all three derivative branches (“progenitors”; **Fig. 3B)**. All three lineages increasingly express the transcriptional HLH inhibitor *ID4*, as *SoxC* diminishes.

After gastrulation, many additional cell types differentiate, including multiple gland, neuronal, and secretory cells. These additional cell types are represented within the post-gastrula developmental subset as additional trajectories arising from a *SoxC*+ NPC population (**Fig. 3C, E**). Altogether, these data indicate a stable presence of NPCs until the primary polyp stage (**Fig. 3C, D**) and throughout homeostasis in the adult polyp **(Fig. 3E, F)**. Consistent with the relationships at the gastrula stage, we observed differentiation trajectories for cnidocytes (e.g. *ZNF845*+, **Fig. 3C, E**) and neurons (e.g. *AshA+, Nanos1*+, **Fig. 3C, E**), while the origin of the mucin gland cells is detectable only by expression of *Not-likeC* (**Fig. 3D, F**). Similar to mucin+ gland cells (“gland.M”), tentacle retractor muscle cells marked specifically by the bHLH transcription factor gene *Nem64* (a recently uncovered master regulator of tentacle retractor muscle development (**Fig. 3D**,**F**, (Cole et al., 2020)), fall within a distinct partition, indicating that we did not capture any intermediate cells and somewhat masking their developmental origin. However, a small cluster of early progenitors is evident in both the developmental and tissue subsets (origin red dashed trajectory **Fig. 3C**,**E**). This is corroborated by expression of the NPC gene sets in the retractor muscle, as well as detectable *Nem64* expression within the NPCs (**Fig. 3 D**,**F**). Trajectories leading to digestive gland cells (*AshC*+) and additional sensory-secretory cells (“S1”, “S2”) appear, as well as an expansion of the neural populations indicative of an expanding neurosecretory repertoire (**Fig. 3C**,**E)**. Apart from the main neuronal trajectory, we identified an independent neuronal pathway arising from the NPC population which terminates as a single neuronal population (“eN.1”, **Fig. 3C, E**). This molecular pathway has a distinct signature which consists of the cnidarian Beta3/Oligo/Mist-like family bHLH transcription factor *Nem5* and the zinc finger transcription factor *PRD14-like3* (**Fig. 3D, F, Fig. S3J**).

We note that the expression pattern of the zinc finger transcription factor *Insulinoma-1* (*Insm1)* clearly identifies a stable subdivision within our neurosecretory partition (**Fig. 3D, F**). *Insm1* expression unites the majority of neurons (“Ni+”) with the expansion of sensory-secretory neuroglandular populations (“S1”, “S2”, “gland.D”, “gland.M”). The absence of *Insm1* in a subset of neurons (“Ni-”) is accompanied by expression of *Pou4* (**Fig. 3C, E**), previously described to drive the terminal differentiation of ectodermal sensory cells and endomesodermal neurons (Tournière et al., 2020). The *Insm1-/Pou4+* expression domain also expresses the nuclear receptor *tailless* (*TLL-like*) in a subset of “Ni-” neurons and in the S2 lineage (**Fig. 3D, F**), and includes the previously mentioned neuronal population “eN.1” and a population of hair cells (**Fig. S3V-W**, (Ozment et al., 2021)).

Within the adult tissues subset, three additional cell clusters appear: an uncharacterized cluster (S3) with no detectable relationships to other clusters, as well as primary germ cells (PGCs, **Fig. S3G**) and early oocytes (**Fig. 3E**). The PGCs are found to emerge first from the cycling cells in polyp libraries, followed by the establishment of the NPC transcriptomic profile. Similar to the developmental subsets (**Fig. 3A**), NPCs are predicted to derive from proliferative precursors, which then enter mitotic arrest as detected by the expression of a cyclin-dependent kinase inhibitor (*CDN1A-like*; **Fig. 3B, D, F**). This cycling precursor population is analyzed in more detail elsewhere (Denner et al., in preparation). In summary, the reconstructed trajectories are similar and robust between all three developmental subsets. Thus, the molecular mechanisms of neurosecretory cell differentiation are maintained from embryogenesis until adulthood, suggesting that embryonic /larval processes also ensure homeostasis during later stages.

### *Nanos1* marks sensory and ganglion neurons in both cell layers

Our comprehensive characterization of the neurosecretory trajectories across all stages predicted an early role in neuronal specification for the mRNA-binding protein *Nanos1* (**Fig. 4A)**, one of the two cnidarian *Nanos* paralogs in *Nematostella* (**Fig. S3D, *Fig. S4, Fig. S5***, *Extavour et al., 2005). Nanos* codes for an RNA-binding protein involved in translational repression, which is well known for its role in the germline and stem cells (Kang et al., 2002; Köprunner et al., 2001; Subramaniam & Seydoux, 1999; Tsuda et al., 2003; Wang & Lehmann, 1991) but also in axial patterning of the early *Drosophila* embryo (Curtis et al., 1995). To validate the putative somatic role in neurogenesis, we generated a transgenic reporter line under the control of the *Nanos1* promoter. In line with the single cell RNAseq data, this transgenic line showed fluorophore expression in numerous sensory and ganglion neurons in both cell layers (**Fig. 4A-i**). We also dissociated adult animals and identified mCherry expression in undifferentiated cells and ganglion neurons in cell spreads **(Fig. 4A-iii)**, which confirmed the predicted role of *Nanos1* in neurogenesis.

**Figure 4:**
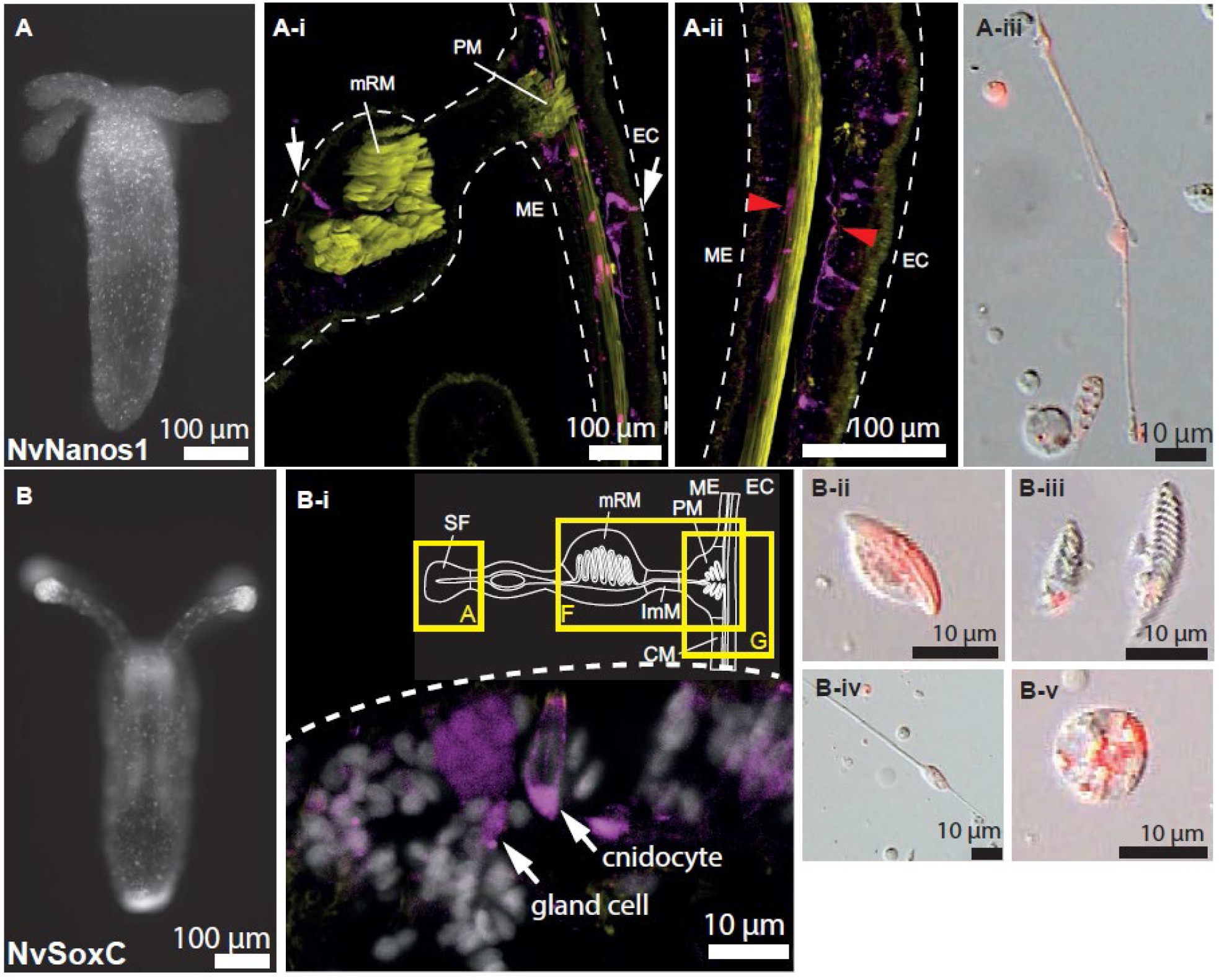
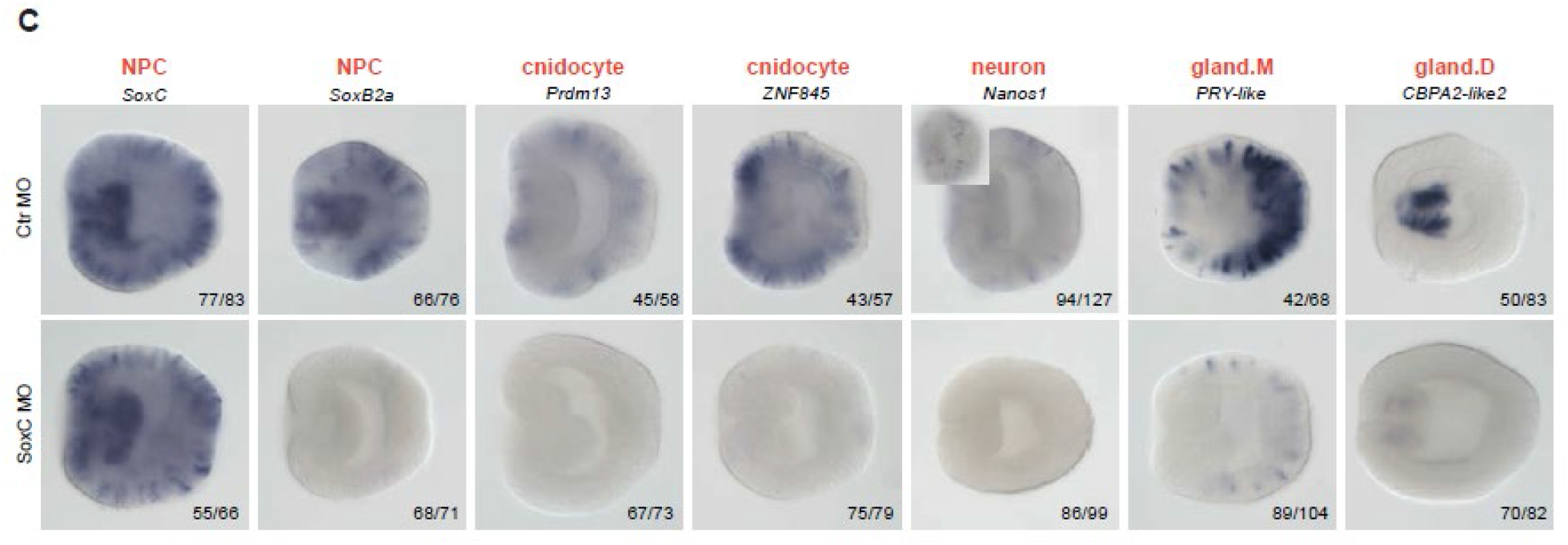
SoxC acts upstream of SoxB2a in the specification of neural progenitor cells. A) Reporter protein expression in the Nanos1::mCh line in a primary polyp. Neurons in ecto-and endomesoderm (A-i, A-ii, sensory neurons indicated by white arrows, ganglionic neurons indicated by red arrowheads), a NPC and a ganglionic neuron (A-iii). B) Characterization of the SoxC::mCh line. Single-cell expression in a primary polyp (B), gland cells and nematocytes in the septal filament (B-i), and in dissociated cells. Reporter protein expression in a nematocyte (B-ii), neuron (B-iii), spirocytes (B-iv), and putative gland cell (B-v). C) Functional validation of SoxC as a key specification factor of the neurosecretory progenitor population. Knockdown of SoxC diminishes the expression of the NPC marker SoxB2a and the derivatives of the NPC population, cnidocytes, neurons, and gland cells at 2dpf.

### SoxC is a key regulator of the entire neurosecretory lineage

The broad expression of HMG transcription factor *SoxC* within both the cycling precursors, the multipotent NPCs, as well as extending along all trajectories, indicates it is a potential key regulator of the neurosecretory lineage (**Fig. 3B, D, F, SI_1**). To trace the fate of *SoxC-*positive cells throughout the life cycle experimentally, we generated a *SoxC::mCherry* transgenic line (**Fig. 4C, Fig. S5**). We observed mCherry+ cnidocytes in ectodermal tissues as well as gland cells in the septal filament and endomesoderm (**Fig. 4C-i**). In cell suspensions generated from transgenic juvenile polyps, a diversity of derivatives of the NPC populations were identifiable by fluorophore expression, including neurons, gland cells and cnidocytes, corroborating the predictions of our sc-transcriptomic data **(Fig. 4C-ii-v)**. Next, we wished to test the function of *SoxC* in the specification of the whole neurosecretory complement. We therefore knocked down *SoxC* using an antisense morpholino (**Fig. S2C**). Indeed, *SoxC* knockdown depleted cell type-specific marker genes of cnidocytes (*ZNF845, Prdm13*), gland cells (*PRY-like, CBPA2-like2*) and neurons (*SoxB2a, Nanos1*) (**Fig. 4D**). Notably, the abolished expression of well-characterized NPC determinant *SoxB2a* (Richards & Rentzsch, 2014) confirms the prediction that this gene is downstream of *SoxC*. Thus, we identified *SoxC* as the most upstream regulator currently identified in the specification of neurosecretory progenitor cells, which give rise to all cell types constituting the neurosecretory partition.

Our predicted differentiation trajectories throughout the life cycle lay the foundations for the construction of a network model of the specification of the neurosecretory lineage **(Fig. 5**). We identify a population of undifferentiated cycling cells, which either become primordial germ cells or adopt the neurosecretory fate, specified by *SoxC* **(Fig. 5**). Primary cell fate decisions during embryogenesis (shaded grey pathways) drive the formation of cnidocytes, neurons and mucus-producing gland cells **(Fig. 5**). Subsequently, additional cell types such as digestive gland cells and the tentacle retractor muscle are generated by the common progenitor population, following stable differentiation trajectories throughout the life cycle (**Fig. 5**).

**Figure 5:**
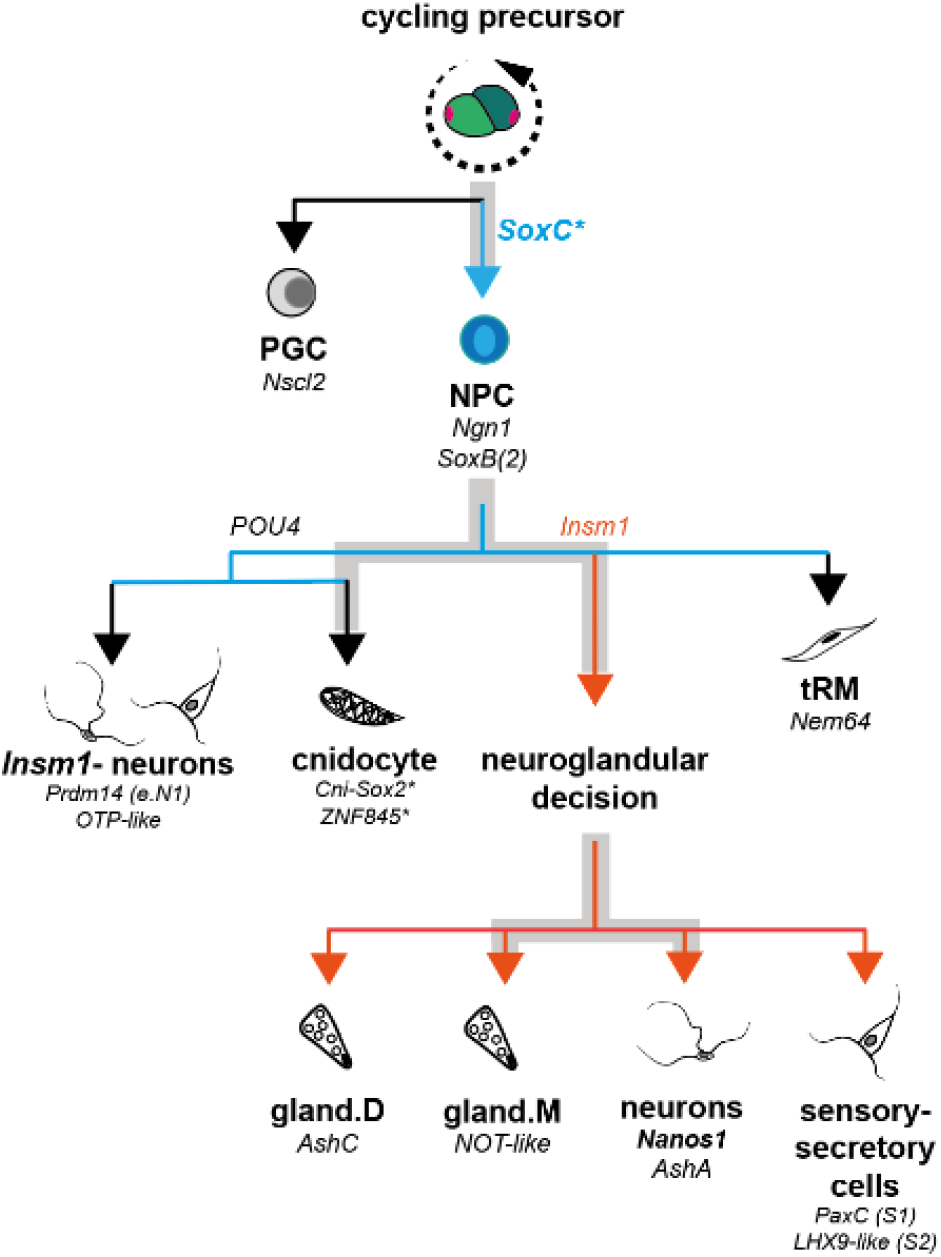
Molecular decision making underlying the population structure of neurosecretory lineages predicted by our dataset. Grey shading indicates decisions made during embryogenesis which persist during homeostasis. Asterisks mark transgenic lines characterized in the scope of this paper.

## DISCUSSION

Neurons, cnidocytes and gland cells represent the major differentiation products in cnidarians (David, 2012; Fujisawa & David, 1981; Galliot & Quiquand, 2011; Hager & David, 1997; Lindgens et al., 2004; Technau & Holstein, 1996). Yet, there is still relatively little known about the molecular and genetic cascades governing these differentiation trajectories. In this study, we report a developmental single cell transcriptomic map of cell type specification in the sea anemone *Nematostella vectensis* with a focus on the neurosecretory cell complement. The high resolution of our dataset allows the reconstruction of differentiation trajectories, unfolding the population structure of all neurosecretory lineages of *Nematostella* and deepening our understanding of the relationships between the previously reported complement of cell types (Sebé-Pedrós et al., 2018) and their common developmental origin in a multipotent progenitor population.

### Cnidocyte trajectory shows unusual diverging and converging transcriptome profiles

Our reconstructed cnidocyte trajectory predicts several upstream regulators of cnidogenesis such as *Sox2*, which is a member of a *Nematostella* specific group of *Sox* genes (and not a homolog of vertebrate *Sox2*, (Babonis, Enjolras, Reft, et al., 2021; Magie et al., 2005; Schwaiger et al., 2021). *Sox2* is coexpressed with other transcription factors such as *ZNF845, Myc4*, and *PRDM13*. ZNF845, belonging to the large family of Krüppel-like zinc finger transcription factors like the pluripotency determinant Klf-4, represents a member of a group of zinc finger genes branching basally to all other Klf families, which has led recently to the proposal that the evolution of this novel cell type may be facilitated by lineage-specific gene duplications (Babonis, Enjolras, Ryan, et al., 2021). This regulatory module is indicative of an early primed cnidocyte state, which is validated by our transgenic lines. Decision-making between distinct subtypes occurs downstream of this transcriptomic state, and is accompanied by subtype-specific transcription factors and effector genes, such as the late differentiation marker *CP1*, a cnidocyte-specific peptidase, which is excluded from the spirocytes.

Cnidocytes are an evolutionary novelty of cnidarians and the phylum-defining trait, and different cnidarian classes have evolved several subtypes, which differ mainly by their capsule type but also by their toxin complement. Besides the three morphologically described subtypes (Zenkert et al., 2011), we also identified two additional types: a larva-specific cnidocyte and an unknown type, which only expresses *Nematostella*-specific genes. This shows that the cnidocyte complement is more diverse than previously appreciated. All subtypes of cnidocytes show distinct trajectories and gene batteries, presumably required to produce the different capsule types. Remarkably, in the final step of differentiation, mRNA for capsule-forming genes is diminished and all subtypes converge towards a common transcriptome profile of the mature cnidocyte ready for discharge. This divergence-convergence pattern of the transcriptomic profile is to our knowledge unique and can only be explained by a transcriptional downregulation of the genes responsible for the specific capsule subtypes, followed by the activation of a common maturation profile. Convergence of structurally distinct subtypes can also be identified in the single cell transcriptome data from *Hydra* (Siebert et al., 2019) suggesting conserved molecular underpinnings of the distinct phases of cnidogenesis in anthozoans and hydrozoans.

#### Neurons and gland cells have a common regulatory signature

While cnidocytes and neurons have commonly been regarded as sister cell types (Richards & Rentzsch, 2014), our data suggest that neurons and gland cells are actually more closely related and have a common progenitor cell, characterized by *Insm1* expression, which supports recent evidence of this relationship (Tournière et al., 2021). This also reveals similarities to the relationships within the interstitial cell lineage in *Hydra*, where neurons and gland cells have a common progenitor state, while nematogenesis is distinct (Siebert et al., 2019). The question arises, whether the common progenitor cell population for neurons and gland cells is purely reflective of the developmental relationship, or whether this also indicates an evolutionary sister cell type relationship. Indeed, there is evidence from several other animal phyla that support the idea of an ancestral sister cell type relationship of neurons and gland cells. Neurosecretory cells in the forebrains of both vertebrates and invertebrates integrate sensory functions and neuropeptide secretion (reviewed by Hartenstein, 2006; Vigh et al., 2004). Furthermore, they have a conserved regulatory code, which further supports the secretory character of ancestral neurons (Tessmar-Raible, 2007; Tessmar-Raible et al., 2007). Moreover, there are striking similarities in the molecular profile and cellular behavior of motor neurons of the ventral nerve cord and secretory pancreatic cells, including a number of shared transcription factors (e.g. *islet, pax6, mnx, nk6 and hnf6*) (Arendt, 2021). Notably, we recently detected a similar shared profile of pharyngeal cells giving rise to secretory digestive and insulinergic gland cells and neurons in *Nematostella*, suggesting an ancestral origin in sensory-neurosecretory cells of the mucociliary sole (Steinmetz et al., 2017). This pharyngeal tissue and its cellular derivatives, including digestive gland cells, insulin-peptide secreting cells, and a number of neurons, also express *FoxA* (Steinmetz et al., 2017). Interestingly, *FoxA2*, which is an endodermal marker in vertebrates, is also expressed in the notochord and the floor plate of the neural tube, where it contributes to the specification of specific dopaminergic neurons (Ang, 2009). Thus, there are multiple common regulators of neuronal and gland cells in sea anemone and vertebrates.

#### SoxC acts as a conserved upstream regulator of all neurosecretory lineages

We newly identified *SoxC* as a key determinant of the entire neurosecretory partition. The *SoxC* transgenic line shows small cells with stem cell appearance as well as neurons, cnidocytes and gland cells in both cell layers (**Fig. 4B**). Knockdown of *SoxC* abolishes marker genes for these cell types including the expression of *SoxB2a*, indicating that *SoxC* acts upstream of *SoxB2a. SoxB2a* is a key determinant of neural progenitor cells in *Nematostella* and this role is highly conserved in bilaterians (Bylund et al., 2003; Graham et al., 2003; Richards & Rentzsch, 2014). *SoxC* is also transiently expressed in the interstitial stem cell differentiation trajectory in *Hydra*, marking the onset of differentiation into neurons and nematoblasts (Siebert et al., 2019), consistent with a conserved role of *SoxC* in the differentiation pathways of cnidarian progenitor cells. So far, a role of *SoxC* in gland cell differentiation has not been documented in *Hydra*, but it is known that interstitial stem cells also give rise to gland cells (Schmidt & David, 1986; Siebert et al., 2019). The role of SoxB and SoxC transcription factors specifying the neurosecretory lineages of *Nematostella* reveals interesting parallels to the vertebrate neural crest, which undergoes differentiation into diverse cell types such as the peripheral nervous system, pigment cells, bone and cartilage (Bhattaram et al., 2010; Hong & Saint-Jeannet, 2005; Tang & Bronner, 2020). Interestingly, neural crest cells also contribute to secretory cells by invading the developing pituitary gland, where they give rise to hormone-secreting cells and pericytes (Ueharu et al., 2017). Consistent with an essential role of SoxC in neural crest specification, knockdown of *SoxC* genes in *Xenopus* and lamprey causes defects in neural crest-derived structures (Uy et al., 2015).

Members of the SoxC group act also as cell fate determinants in CNS formation, especially in neural induction and subsequent steps of neuronal differentiation including dendritogenesis, establishment of neuronal projections, migration of precursors and subtype specification (for review see (Kavyanifar et al., 2018)). During early mouse embryogenesis the SoxC family members Sox4 and Sox11 regulate the survival of early neural precursors and also promote neuronal differentiation (Chen et al., 2015; Cheung et al., 2000). Besides its role in neuronal differentiation, *Sox4* has also shown to have a crucial role in endometrial carcinomas, by transcriptionally activating *Tcf4* and hence the ß-catenin/Tcf4 dependent proliferation (Saegusa et al., 2012). There is also evidence for a role of Sox4 in the development of sensory-secretory cells in vertebrates, e.g. in tuft cells in the thymus and intestine (Gracz et al., 2018; Mino et al., 2022). While the *SoxC* homolog does not seem to play a role in neurogenesis in *Drosophila*, in the annelid *Platynereis dumerilii* and the spider *Parasteatoda tepidariorum, soxC* homologs are expressed in the neurogenic field of the developing embryo larva (Kerner et al., 2009; Paese et al., 2018). In both vertebrates and the sea anemone *SoxC* expression precedes *Soxb2a* and *Neurogenin* expression, however, they also extensively overlap during neurogenesis and at least in vertebrates appear to form distinct complexes involved in the definition of neuronal subtypes (Chen et al., 2015). This indicates an ancestral function of *Sox* genes in key lineage decisions in multipotent neural progenitors, pre-dating the evolution of complex nervous systems.

#### Neurosecretory differentiation trajectories are maintained after embryogenesis

The specification of cell types and their homeostatic maintenance and replacement by stem cells are a major interest of developmental biology. Notably, in vertebrates and many other bilaterians, the central nervous system develops during embryonic and early developmental stages, but there is little neurogenesis at later stages (Kempermann et al., 2018; Ming & Song, 2011). As a consequence, the central nervous system shows restricted regenerative capabilities in cases of damage or cell death. Cnidaria are of particular interest in this regard, because they have remarkably long life spans, with stunning plasticity in body size regulation and their cells have to be continuously replaced (Watanabe et al., 2009). We identified a common neurosecretory trajectory emerging from a multipotent neural progenitor cell (NPC) population and show that it is present throughout the life cycle. Our data suggest that the same molecular hierarchies and pathways that are established during early embryogenesis continue to be active throughout the life of the polyps allowing for a constant replacement of neurosecretory cells. The molecular regulation of neurogenesis is largely conserved between cnidarians and bilaterians (Rentzsch et al., 2017). A mechanistic understanding of how homeostasis is regulated in cnidarians provides future opportunities to address the limited capacity for regeneration in the central nervous system of adult vertebrates.

## Supporting information

SI_3. Gene model annotation list

SI_1. Related to Figure 1. 3d-pro

SI_2. Related to Figure 1 and 2. Differential marker gene expression lists

## Acknowledgements

We thank Elly Tanaka, Sasha Mendjan and Florian Raible for critical reading of the manuscript and the Technau group for continuous discussions. We thank Julia Hagauer and Oliver Link for collecting samples and assistance with library preparation. We thank Angela Caballero Alfonso, Wolfgang Göschl and Vendula Stejskalova for providing excellent care of the Nematostella facility. Sequencing was performed at Vienna BioCenter Core Facilities (VBCF) and Novogene. We are grateful to the support by the CIUS imaging facility of the University of Vienna. This work was supported by grants of the Austrian Science Fund FWF to U.T. (Grant DOC 72 doc.fund), A.G.C. (P31018-B29), and G.G. (P30404-B29 and P32705-B) as well as the Research platform SINCEREST to U.T. funded by the University of Vienna.

## Author contributions

Conceptualization, AGC, AD, JS, and UT; Formal Analysis, AGC and JS; Investigation, JS, AGC, AD, TL, AR, RR, ET, ML; Writing - Original Draft, AGC, AD, JS; Writing - Review & Editing, AGC, GG, UT; Supervision, AGC, GG, and UT; Funding Acquisition, AGC, GG and UT

## METHODS

### *Nematostella* culture

Adult polyps were cultured in *Nematostella* medium (“NM”, 1/3 artificial sea water) at 18°C in the dark. Spawning was induced by exposure to light and increased temperature (Fritzenwanker & Technau, 2002). Fertilized egg packages were treated with a 3% cysteine/NM solution to remove the egg jelly. Embryos were raised at 21°C until collection for cell dissociation.

### Generation of single-cell suspensions

Animals of all stages were selected based on their morphology, pooled in pre-coated 0,5 ml Eppendorf tubes and washed in NM. Enzymatic dissociation was performed with 10X TrypLE Select (Gibco™, catalog number A1217701). Enzyme concentration and dissociation time were adapted to the individual developmental stages. Dissociation was stopped by adding 1%BSA/PBS. Three washing steps were performed before resuspending the cell pellet in BSA/PBS. The cell suspension was kept on ice until further processing. For assaying cell viability and quantifying the cell concentration we used a Cellometer X2 (Nexcelom), setting the viability threshold to a minimum of 80%. Cell suspensions were then diluted to 1000 cells/uL prior to loading into the 10xGenomics platform.

### Cryopreservation

The cell suspension was generated as described above for 4d planula larvae, but then pelleted and carefully resuspended in pre-chilled 0,5% FBS in NM and supplemented with DMSO to a final concentration of 10%, according to (Wohnhaas et al., 2019). The cells were then transferred into cryo vials, placed in a CoolCell (BioCision) and stored at -80°C for 4 days. For recovery, cells were picked up from the -80°C freezer with a pre-cooled -20°C thermo block and then incubated at 20°C for 6-8 min. FBS was supplemented and cells were washed two times, resuspended in BSA/NM and then processed through quality control and quantification like all other live cell suspensions. The resultant library was similar in cell content as the live-captured control.

### Single-cell RNA sequencing

Single-cell suspensions were loaded into the 10X Genomics platform for encapsulation. Sequencing libraries were prepared according to the standard 10X protocol, the version of the respective 10X chemistry is noted in **Fig. S2**. Illumina sequencing was performed at the NGS facility of the Vienna BioCenter Core Facilities or Novogene. Raw sequencing data was processed with the CellRanger 2.1.0 pipeline, using default parameters and forcing a recovery of 7000 cells for each library.

### Reference transcriptome

For mapping the reads we used a customized transcriptome (M. S. David Fredman, Fabian Rentzsch, and Ulrich Technau). (figshare,2013) /https://figshare.com/articles/dataset/Nematostella_vectensis_transcriptome_and_gene_models_v2_0/807696). To increase mapping efficiency, gene models were extended by 1000 bp in the 3’ direction, or until the start of the next gene model in the same orientation.

### Single cell transcriptomic analysis of single libraries

Count matrices were imported into R and processed with the R-package Seurat vs3 and vs4 (Stuart et al., 2019). In order to retain only high-quality cells for downstream analysis we applied the following filtering criteria: Cell multiplets were removed by using a library-specific cutoff for aberrantly high UMI counts, as specified in Table S1. Low-quality cells were excluded by setting the percentage of mitochondrial counts to a maximum of 10% and the minimal gene expression to 250 or 300 genes/cell depending on the 10x chemistry used. After this initial quality control step we normalized our data with Seurats default “LogNormalize” function and identified the 2000 most variable genes with the “FindVariableGenes” function with default parameters. Scaling was performed with the “ScaleData” function, using the variable features. Informative principle components were chosen as those with the highest standard deviation and used as input for the clustering analysis, while PCs that show a plateau in their SD were excluded. Dimensional reduction was performed with UMAP using the same PCs as in the clustering. Differentially expressed genes were identified with the FindAllMarkers function. The ranked lists of putative cluster markers were analyzed to determine cluster identities and validated with the FeaturePlot function. For visualization of gene expression values, missing data was inferred using Markov Affinity-based Graph Imputation of Cells (MAGIC) with the R package Rmagic (vs. 2.0.3) (Van Dijk et al., 2018).

### Merging all single libraries

The raw UMI-reduced data matrices from each individual dataset were concatenated, and a corresponding Seurat object was generated and processed as above with the following exceptions: first the top 1000 variable features were selected from each individual library, and the combined set was imported into the merged object. Normalized counts were then scaled within each library prior to calculating the principal components. We find that this approach allows for biologically relevant data integration and ameliorates most batch effects while maintaining developmental signals within the data.

### Subset analyses

Cell lineages of interest were extracted from the merged dataset and processed through the same pipeline as single libraries, with the only deviating parameter that the full dataset scaling was imported. For generation of cell trajectories, the Seurat subset object was converted to a cell data object and used with Monocle3 vs1.0.0 (Cao et al., 2019; Levine et al., 2015; Qiu et al., 2017; Traag et al., 2019; Trapnell et al., 2014), partitions and cell clusters were calculated with a resolution parameter of 0.01, and the trajectory was calculated with the learn_graph function. Pseudotime was estimated by manually selecting the progenitor population as the starting point. To estimate the differentiation state, the package cytoTRACE vs0.1.0 (Gulati et al., 2020) was run on the raw count matrix.

### Generation of transgenic lines

The transgenic lines were generated as described (Renfer et al., 2010). In short, we cloned putative promoter (*NvSoxC*: 3603 bp, *NvSox2*: 3134 bp, *NvMMP1*: 1959 bp, *NvNanos1*: 1525 bp) and enhancer regions (*NvSox2*: 2834 bp, *NvNanos1*: 2863 bp) upstream and downstream of the target genes defined by previously identified epigenetic signatures (Schwaiger et al., 2014). Identified sequences were cloned in front of the open reading frame of either mCherry or mOrange fluorophores on a transgenesis vector based on the p-CRII-TOPO backbone (Shaner et al., 2004). The *I-SceI*-digested transgenesis plasmids were injected into fertilized eggs at a concentration of 25 ng/µl. Animals were raised and crossed to wild-type polyps to generate F1 heterozygous transgenics.

### Gene knockdown

For SoxC knockdown we microinjected the translation-blocking morpholino oligonucleotide GTAGCACCATCATCATCACTAGAA (500 ng/µl). The activity of the morpholino was tested by co-injecting it with mCherry mRNA fused in-frame at its 5’ end with either the wild-type or the 5-mismatch recognition sequence of the morpholino recognition sequence **(Fig. S2**). As a control, the previously described control morpholino oligonucleotide (Kraus et al., 2016) was used. Morpholinos were injected into zygotes as previously described (Lebedeva et al., 2021).

### *In situ* hybridization

For whole-mount in situ hybridization (ISH), the embryos were fixed with 4% PFA/PBS for 1 hour at room temperature. Then, fixed embryos were washed with PTw (PBS, 0.1% Tween 20) 3 times for 5 minutes, transferred into 100% methanol, and stored at -20°C. ISH was performed as described previously (Kraus et al., 2016) with minor changes in the RNA probe detection step: anti-Digoxigenin-AP Fab fragments (Roche) diluted 1:4000 in 0.5% blocking reagent (Roche) in 1× MAB were used. After overnight incubation in anti-Digoxigenin-AP Fab fragments at +4°C, embryos were washed with PTw 10 times for 10 minutes, rinsed 2 times for 5 minutes with alkaline phosphatase buffer and stained with NBT/BCIP (Roche) as described previously (Kraus et al., 2016). After clearing with 86% Glycerol, embryos were imaged with a Nikon 80i compound microscope equipped with the Nikon DS-Fi1 camera.

### Data availability

Raw sequence data and corresponding count matrices have been submitted to GEO (https://www.ncbi.nlm.nih.gov/geo/) and are currently being processed (reference to be provided when available). Low-dimensional cell embeddings from each analysis are hosted on the USCS Cell Browser for user-specified gene expression exploration (https://cells.ucsc.edu/), and reviewer access to this platform can also be provided upon request. The code for generating results is available upon request.

## SUPPLEMENTAL FIGURES

**Figure S1: Cell type diversity within individual single-stage- and tissue-specific libraries**. Color code corresponds to the 12 coarse clusters in Fig. 1C’. Abbreviations: AO: Apical organ, gland.D: Digestive gland cell, gland.M: Mucus-producing gland cell, iMM: intermuscular membrane, N: neuronal, NPC: Neural progenitor cell, S: sensory-secretory. *indicates tissues that were dissected from the same polyp.

**Figure S2A: Specifications of all individual libraries that compose the merged dataset. Figure S2B: Published marker genes used for cluster annotation**.

**Figure S2C: Testing the efficiency of the SoxC-morpholino**. Upon co-injection, SoxC MO inhibits the translation of the mCherry mRNA fused in-frame to the wild type morpholino recognition sequence. Co-injection of the SoxCMO with mCherry mRNA carrying a 5-mismatch recognition sequence for the SoxCMO is not repressed.

**Figure S3: Expression patterns of surveyed marker gene candidates**. V-W) Insm1 and POU4 expression patterns denote a clear subdivision. V) UMAP projection of neurosecretory cell types, excluding cnidocytes. The main subgroups include neurons, gland cells and sensory-secretory groups S1-S3. W) The Insm-negative/POU4-positive domain includes distinct cell populations such as endomesodermal neurons and hair cells. The hair cell marker is polycystin 1 (Ozment et al 2021).

**Figure S4: Bayesian consensus tree of Nanos proteins based on amino acid alignment, indicating independent duplications of a single ancestral Nanos gene in the Bilaterians and Cnidarians**. Species abbreviations: Dre: Danio rerio, Hsa: Homo sapiens, Mmu: Mus musculus, Xla: Xenopus laevis, Bfl: Branchiostoma floridae, Dme: Drosophila melanogaster, Hro: Helobdella robusta, Hvu: Hydra vulgaris, Che: Clytia hemisphaerica, Hec: Hydractinia echinata, Pca: Podocoryne carnea, Nve: Nematostella vectensis, Emu: Ephidatia muelleri. Vertebrata are displayed in green, Arthopoda in yellow, Annelida in blue, Cnidaria in pink and porifera in red. Numbers next to branches indicate bootstrap support (1000 iterations).

**Figure S5: Reporter protein expression in Sox2::mOrange, CP1::mCherry, Nanos1::mCherry and SoxC::mCherry transgenic lines**. A) Sox2 expression in a cross-section of a tentacle. Magnified nematocytes with basally located cytoplasm and nucleus (green arrowheads) are outlined. B) Detail picture of Sox2+ cnidocytes in a tentacle. Spirocytes are indicated by white arrows, nematocytes by green arrowheads. C) Brightfield and fluorescence picture of a tentacle showing CP1+ nematocytes. Reduced dot-like cytoplasm is indicated by green arrowheads in selected cases. Non-fluorescent spirocytes are outlined in selected cases. D) Nanos1+ sensory neurons in ectoderm (blue arrowheads) and endomesoderm (red arrowheads). E) Distribution of Nanos1+ neurons in a cross-section of a juvenile polyp. Neurons in the septal filament and endomesoderm surrounding the retractor muscle are indicated by white arrows. F) Distribution of Nanos1+ neurons in a longitudinal section of a juvenile polyp. G) SoxC expression in a cross-section of a tentacle. Spirocytes and a nematocyte with basally concentrated cytoplasm (green arrowhead) are indicated in the inlet. H) Detail picture of SoxC+ cnidocytes in the body column (yellow arrowheads). I) SoxC+ gland cell (white arrow) in the vicinity of the parietal muscle. Fluorescent reporter proteins displayed in magenta/red, Phalloidin in yellow/green, DAPI in grey/blue. Abbreviations: SF: Septal filament, mRM: Mesentery retractor muscle, ImM: Intra-muscular membrane, PM: Parietal muscle, EM: Endomesoderm, EC: Ectoderm, T: Tentacle, Pha: Pharynx.

## SUPPLEMENTAL ITEMS

SI_1. Related to Figure 1. 3d-projection of our full developmental time course dataset with *SoxC* expression.

SI_2. Related to Figure 1 and 2. Differential marker gene expression lists

SI_3. Gene model annotation list

