## Supplementary figures and images for "Single cell transcriptomics identifies conserved regulators of neurosecretory lineages"

### SI_1. Related to Figure 1. 3d-pro

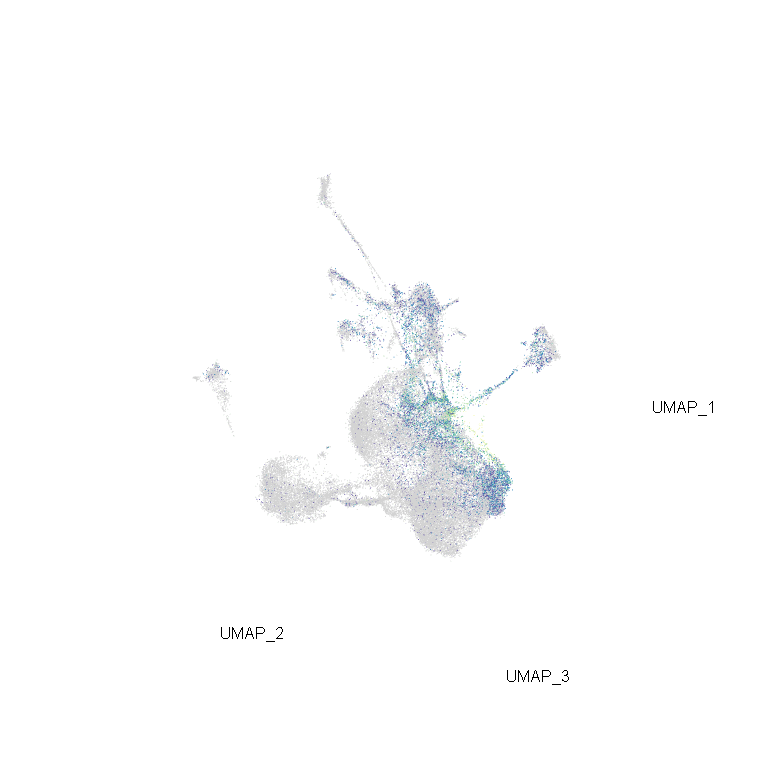
