## Supplemental Figures for "Single cell transcriptomics identifies conserved regulators of neurosecretory lineages"

**Figure S1. Cell type diversity within individual single-stage- and tissue-specific libraries.** Color code corresponds to the 12 coarse clusters in Fig. 1C‘. Abbreviations: AO: Apical organ, gland.D: Digestive gland cell, gland.M: Mucus-producing gland cell, iMM: intermuscular membrane, N: neuronal, NPC: Neural progenitor cell, S: sensory-secretory. *indicates tissues that were dissected from the same polyp.

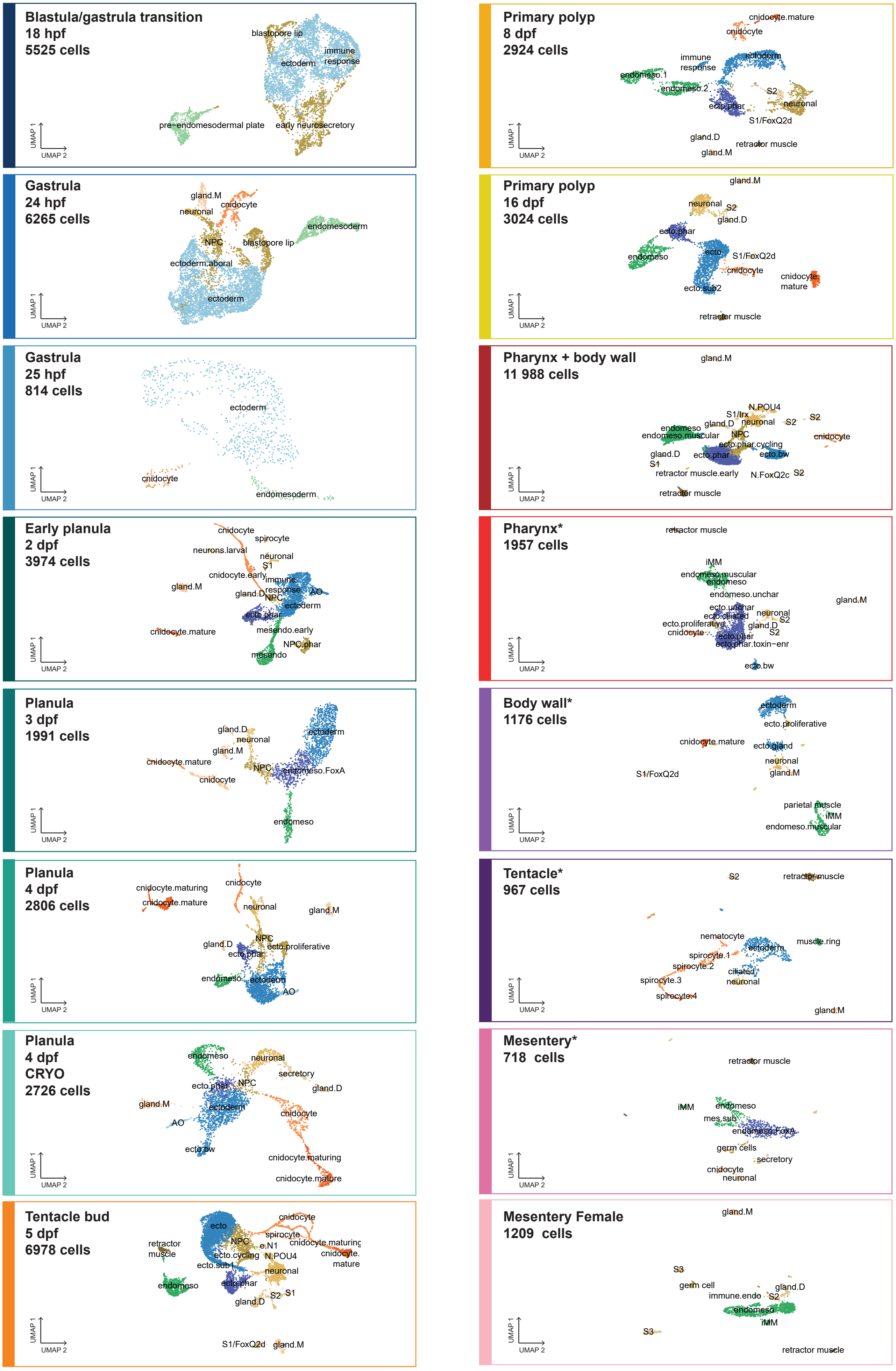

**Figure S2A. Specifications of all individual libraries that compose the merged dataset.**

| | LIBRARY PREPARATION | | | SEQUENCING | CELL SELECTION | | | | | GENE MODEL DETECTION | |
|---|---|---|---|---|---|---|---|---|---|---|---|
| | Developmental stage | Time point (21°C) | 10x chemistry | Saturation | Cell contribution | Min nGene | Max nUMI | Median nGene | Median nUMI | total models | % |
| 1 | Blastula-gastrula transition | 18 hpf | *v3 | 32,7% | 5525 | 300 | 20 000 | 2140 | 5265 | 21982 | 85 |
| 2 | Gastrula | 24 hpf | v2 | 71,4% | 6265 | 250 | 10 000 | 1200 | 2551 | 19941 | 77 |
| 3 | Gastrula | 25 hpf | v2 | 87,3% | 814 | 250 | 40 000 | 482 | 942 | 17633 | 69 |
| 4 | Early planula | 48 hpf | *v3 | 65,2% | 3974 | 300 | 30 000 | 1562 | 4201 | 22637 | 88 |
| 5 | Midplanula | 3 dpf | v2 | 83,5% | 1991 | 250 | 25 000 | 553 | 1576 | 19058 | 74 |
| 6 | Late planula LIVE | 4 dpf | *v3 | 60,5% | 2806 | 300 | 10 000 | 929 | 2032 | 21835 | 85 |
| 7 | Late planula CRYO | 4 dpf | *v3 | 77,7% | 2726 | 300 | 10 000 | 495 | 869 | 21590 | 84 |
| 8 | Tentacle bud | 5 dpf | v2 | 65,4% | 6978 | 250 | 20 000 | 916 | 2330 | 21509 | 84 |
| 9 | Primary polyp | 8 dpf | v2 | 84,7% | 2924 | 250 | 15 000 | 621 | 1916 | 20471 | 80 |
| 10 | Primary polyp | 16 dpf | v2 | 75,2% | 3024 | 250 | 20 000 | 523 | 1495 | 19268 | 75 |
| 11 | Pharynx/body wall | 2 months | v2 | 61,1% | 11988 | 250 | 10 000 | 530 | 996 | 21447 | 83 |
| 12 | Pharynx | 5 months | v2 | 82,0% | 1957 | 250 | 10 000 | 361 | 715 | 17476 | 68 |
| 13 | Body wall | | v2 | 82,0% | 1176 | 250 | 5 000 | 560 | 1563 | 16015 | 62 |
| 14 | Tentacle | | v2 | 83,0% | 967 | 250 | 10 000 | 450 | 1013 | 16383 | 64 |
| 15 | Mesentery | | v2 | 84,0% | 718 | 250 | 15 000 | 367 | 850 | 15933 | 62 |
| 16 | Mesentery.female | adult | *v3 | 93,1% | 1209 | 300 | 20000 | 342 | 917 | 17786 | 69 |

**Figure S2B. Published marker genes used for cluster annotation.**

| | CLUSTER | ANNOTATION USED HERE | ANNOTATION IN REF | REFERENCE |
|---|---|---|---|---|
| 1 | mesendoderm.embryonic | NvSnailA | | Fritzenwanker 2004 |
| 2 | gastrodermis | NvSnailA | | Fritzenwanker 2004 |
| 3 | retractor muscle | Nve-Tpm2 | | Cole 2020 |
| 4 | ectoderm.embryonic | TXT51-like | NvePtx1 | Columbus-Shenkar 2018 |
| 6 | ectoderm.pharyngeal | NvFoxA | | Fritzenwanker 2004 |
| 7 | NPC | NvSoxB2a | NvSoxB(2) | Richards 2014 |
| | | NvNeurogenin1 | NvAth-like | Richards 2015 |
| | | NvMyc1 | v1g38935 | Sébé-Pedrós 2018 |
| | | NvMyc3 | v1g178332 | Sébé-Pedrós 2018 |
| 8 | neuronal | NvAshA | | Layden 2012 |
| 9 | secretory | NvTrypsinB | | Steinmetz 2017 |
| | | NvFoxQ2d | | Busengdal 2017 |
| | | ADAM9-like3 | hemicentin/thrombospondin | Babonis2016 |
| 10 | gland.mucous | NvMucin | | Steinmetz 2017 |
| 11 | cnidocyte | NvNcol3 | | Zenkert 2011 |
| | | EVA1C-like3 | Nematogalectin/Ngal | Babonis 2017 |
| 12 | cnidocyte.mature | FOS-like | Cnido-Fos1 | Sunagar 2018 |

**Figure S2C. Testing the efficiency of the *SoxC*-morpholino.** Upon co-injection, SoxC MO inhibits the translation of the mCherry mRNA fused in-frame to the wild type morpholino recognition sequence. Co-injection of the SoxCMO with mCherry mRNA carrying a 5-mismatch recognition sequence for the SoxCMO is not repressed.

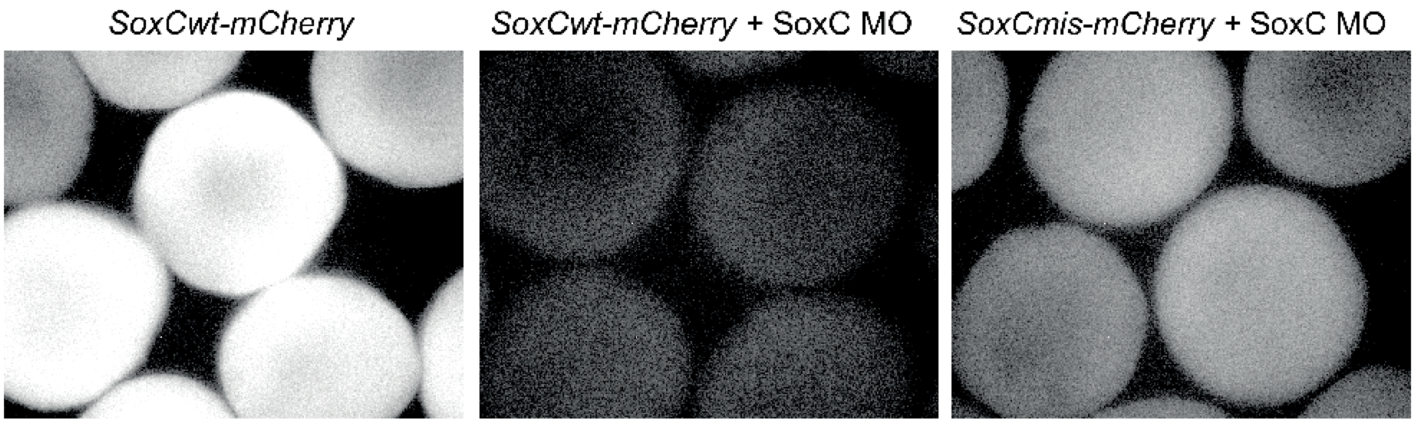

**Figure S3. A-U) Expression patterns of surveyed marker gene candidates. V-W) Insm1 and POU4 expression patterns denote a clear subdivision.** V) UMAP projection of neurosecretory cell types, excluding cnidocytes. The main subgroups include neurons, gland cells and sensory-secretory groups S1-S3. W) The *Insm*-negative/*POU4*-positive domain includes distinct cell populations such as endomesodermal neurons and hair cells. The hair cell marker is polycystin 1 (Ozment et al 2021).

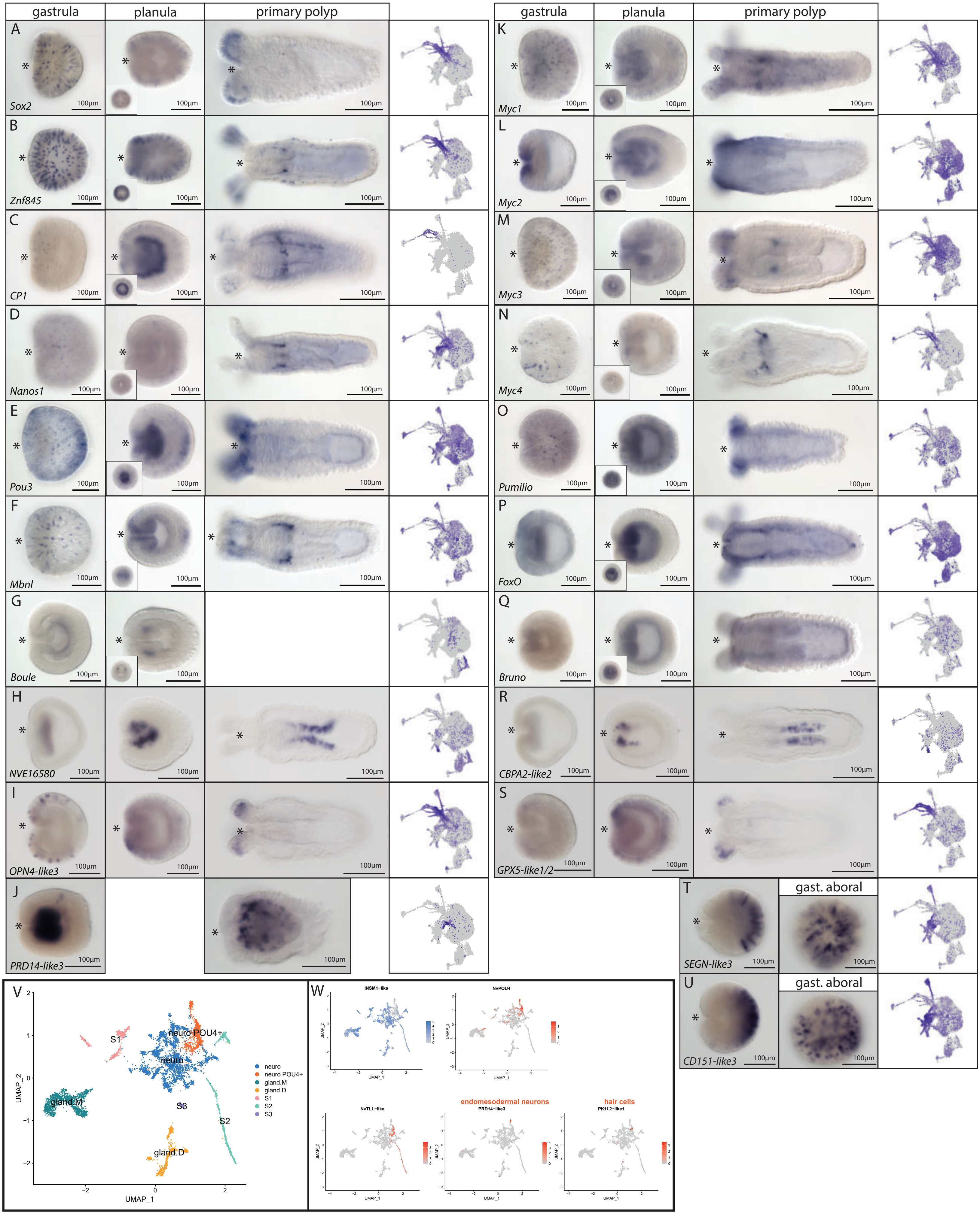

**Figure S4. Bayesian consensus tree of Nanos proteins based on amino acid alignment, indicating independent duplications of a single ancestral Nanos gene in the Bilaterians and Cnidarians.** Species abbreviations: Dre: *Danio rerio*, Hsa: *Homo sapiens*, Mmu: *Mus musculus*, Xla: *Xenopus laevis*, Bfl: *Branchiostoma floridae*, Dme: *Drosophila melanogaster*, Hro: *Helobdella robusta*, Hvu: *Hydra vulgaris*, Che: *Clytia hemisphaerica*, Hec: *Hydractinia echinata*, Pca: *Podocoryne carnea*, Nve: *Nematostella vectensis*, Emu: *Ephidatia muelleri*. Vertebrata are displayed in green, Arthopoda in yellow, Annelida in blue, Cnidaria in pink and porifera in red. Numbers next to branches indicate bootstrap support (1000 iterations).

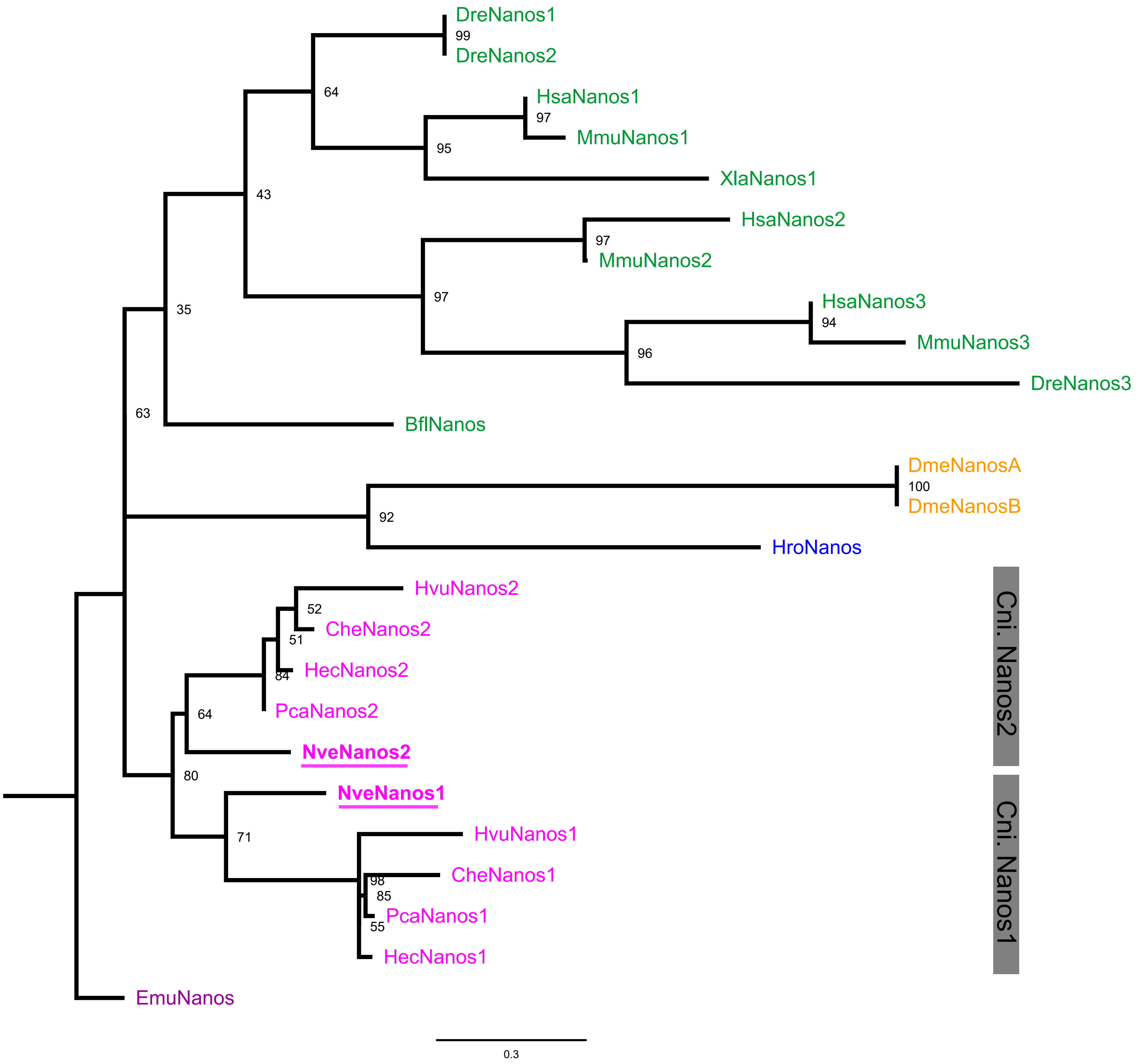

**Figure S5: Reporter protein expression in Sox2::mOrange, CP1::mCherry, Nanos1::mCherry and SoxC::mCherry transgenic lines.** A) Sox2 expression in a cross-section of a tentacle. Magnified nematocytes with basally located cytoplasm and nucleus (green arrowheads) are outlined. B) Detail picture of Sox2+ cnidocytes in a tentacle. Spirocytes are indicated by white arrows, nematocytes by green arrowheads. C) Brightfield and fluorescence picture of a tentacle showing CP1+ nematocytes. Reduced dot-like cytoplasm is indicated by green arrowheads in selected cases. Non-fluorescent spirocytes are outlined in selected cases. D) Nanos1+ sensory neurons in ectoderm (blue arrowheads) and endomesoderm (red arrowheads). E) Distribution of Nanos1+ neurons in a cross-section of a juvenile polyp. Neurons in the septal filament and endomesoderm surrounding the retractor muscle are indicated by white arrows. F) Distribution of Nanos1+ neurons in a longitudinal section of a juvenile polyp. G) SoxC expression in a cross-section of a tentacle. Spirocytes and a nematocyte with basally concentrated cytoplasm (green arrowhead) are indicated in the inlet. H) Detail picture of SoxC+ cnidocytes in the body column (yellow arrowheads). I) SoxC+ gland cell (white arrow) in the vicinity of the parietal muscle.

Fluorescent reporter proteins displayed in magenta/red, Phalloidin in yellow/green, DAPI in grey/blue.

Abbreviations: SF: Septal filament, mRM: Mesentery retractor muscle, ImM: Intra-muscular membrane, PM: Parietal muscle, EM: Endomesoderm, EC: Ectoderm, T: Tentacle, Pha: Pharynx

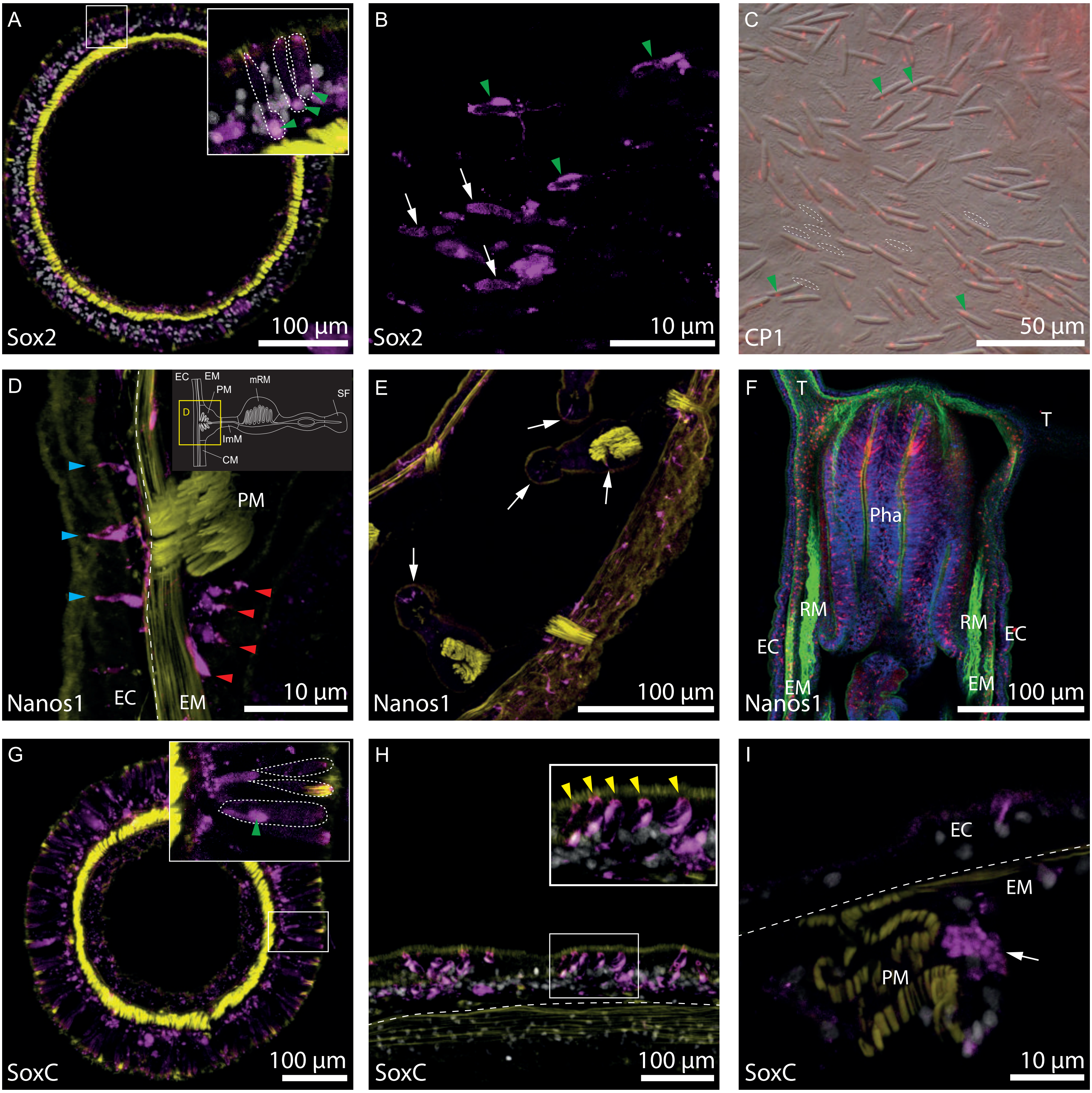
